# Transport of sphingolipids by yeast Npc2 supports phase separation of the vacuole membrane

**DOI:** 10.1101/2025.09.14.676161

**Authors:** Hyesoo Kim, Nicolas-Frédéric Lipp, Israel Juarez-Contreras, Adrian M. Wong, Itay Budin

## Abstract

The yeast vacuole membrane forms ordered microdomains that facilitate micro-lipophagy under nutrient limitation. We previously found that this process involves the intracellular sorting of sphingolipids to the vacuole. While multiple vacuole protein pathways have been identified, corresponding mechanisms for lipid sorting remain undefined. Here we use a range of approaches to identify how endocytic sorting and intraluminal transport of sphingolipids contribute to the formation of vacuole domains. To visualize sphingolipid trafficking, we employed the ceramide analogue BODIPY C12-ceramide (BODIPY-Cer), which is internalized by cells and stains the vacuole. We observed that cells lacking Vps29 and Vps30, proteins involved in endosomal sorting, show altered vacuole domains and accumulate BODIPY-Cer at sites proximal to the plasma membrane. Subsequent incorporation of endocytic-derived ceramide into the vacuole is dependent on the Niemann-Pick Type C 2 protein (Npc2). Loss of Npc2 reduces domain formation and causes BODIPY-Cer to accumulate within the vacuole lumen. Both intra-vacuole trafficking of BODIPY-Cer and membrane phase separation were not dependent on Npc2’s canonical receptor, Ncr1. Lipidomics of isolated vacuoles confirmed that Npc2 independently mediates sphingolipid sorting under micro-lipophagy conditions. In liposome assays, Npc2 robustly transports analogues of ceramide and inositol phosphorylceramide, a complex sphingolipid that is enriched in phase-separated vacuoles. We propose that the enlarged binding cavity of yeast Npc2 is specialized for the incorporation of sphingolipids into the vacuole membrane.

## Introduction

Due to their long, saturated chains and close interactions with sterols, sphingolipids have long been implicated in the formation of ordered membrane domains on cell membranes. We recently observed that sphingolipids are a key lipidic driver for microdomain formation in the yeast vacuole (1), an organelle equivalent to the mammalian lysosome. Vacuole liquid-ordered (Lo) membrane domains form and phase separate from surrounding liquid-disordered (Ld) domains during the transition from exponential growth to stationary growth and are promoted by depletion of nutrients (2). Lo domains play an important role in survival under nutritional restriction, like glucose starvation, because they serve as a platform for lipid droplet docking for micro-lipophagy (3, 4). During micro-lipophagy, neutral lipids like triacylglycerols (TAGs) and sterol esters are hydrolyzed to release fatty acids and sterols to maintain their energy needs and lipid homeostasis (5). Analysis of isolated domain-containing and domain-free vacuoles showed that the former feature increases in sphingolipids, especially inositol phosphorylceramide (IPC). In contrast, whole cell lipidomes were largely unaltered, implicating intracellular sphingolipid sorting. Yeast strains defective in IPC synthesis showed reduced microdomain formation, as well as impaired micro-lipophagy activity (1). These results indicated that trafficking of sphingolipids to the vacuole through unresolved mechanisms is required for domain formation.

Sphingolipid biosynthesis begins in the endoplasmic reticulum (ER) where sphingoid bases are synthesized and *N*-acylated into ceramides. These are then transported to the Golgi apparatus, where their C1 hydroxyl group is modified with polar head groups to generate complex sphingolipids. From the Golgi, these lipids are distributed to other cellular compartments. In yeast, sphingolipid composition varies across organelles: IPC is enriched in Golgi and vacuole, while the plasma membrane (PM) is enriched in mannose inositol phosphorylceramide (MIPC) and mannose di-inositol phosphorylceramide (M(IP)_2_C) (6). Initial steps of ceramide trafficking from ER to Golgi occur via membrane contact sites marked by Nvj2 (7), the ceramide binding protein Svf1 (8), or by vesicular transport (9). Subsequent transport of sphingolipids from the Golgi to PM relies on secretory vesicles, as they fail to reach the PM in the late secretory mutants (10). Sphingolipids are thus likely trafficked to the vacuole primarily through protein trafficking pathways, which rely on vesicular carriers that are routed to the vacuole from other endomembranes.

Vacuole protein sorting (vps) pathways include the carboxypeptidase Y (CPY), alkaline phosphatase (ALP), cytoplasm-to-vacuole targeting (Cvt), endosomal and autophagy pathways (11). The CPY pathway transports both soluble CPY and membrane-bound CPS (carboxypeptidase S) in clathrin-coated vesicles from the late Golgi, through endosomes, to vacuole (12). The sorting of CPY in the Golgi lumen requires its binding receptor Vps10 for proper transport into the vacuole (13). The ALP pathway sends cargo directly from the late Golgi to vacuole, bypassing endosomal compartments. This pathway requires an adaptor protein complex, AP-3 for sorting into non-clathrin coated vesicles, and requires Vps45 for fusion with the vacuole (14). In the endosomal pathway, contents from the plasma membrane and extracellular space are internalized via endocytosis and trafficked through pre-vacuolar compartments and late endosomes. The Cvt pathway is considered a selective autophagic pathway only found in yeast, and delivers specific hydrolases, such as aminopeptidase I or alpha-mannosidase to the vacuole under nutrient-rich conditions (15). Endosomal and autophagic pathways are the two primary routes for intracellular degradation. Late endosomes containing intraluminal vesicles (ILVs) formed by Endosomal Sorting Complex Required for Transport (ESCRT) machinery – thus termed multivesicular bodies (MVBs) – fuse with the vacuole, releasing the ILVs and their cargo into the vacuole lumen for degradation (16). Previously, it was observed that cells lacking Vps4, an AAA-ATPase required for ESCRT-mediated MVB assembly, fail to form vacuole microdomains (2). Autophagic processes include macroautophagy, which requires formation of autophagosomes that fuse with the vacuole, or microautophagy, in which cargoes are directly transported across the vacuole membrane (17). Loss of autophagy in *atg9*Δ cells causes loss of vacuole microdomains (3), suggesting that internalization of cargoes themselves under autophagic conditions might promote domain formation (18).

Both endosomal and autophagic pathways converge at the vacuole for degradation and recycling of biomolecules, ensuring turnover of cellular components and maintaining homeostasis. While this also applies to sphingolipids, it is not known how they are processed inside the vacuole (19). In metazoans, sphingolipids are degraded into their building blocks—sugars, phosphate groups, sphingoid base and fatty acids by multiple enzymes in lysosomes (20), which are functionally equivalent to vacuoles in yeast. Proper degradation and egress of catabolites is critical for maintaining lipid homeostasis, and failure to do so causes lysosomal dysfunction. In humans, defects in sphingolipid degradation cause lysosomal storage diseases (LSDs) that impairs the central nervous system (21). One class of LSDs are Niemann-Pick Type C (NPC) diseases, caused by mutations in two lysosomal lipid transport proteins, NPC1 and NPC2. Both NPC1 and NPC2 are thought to act on lysosomal cholesterol, with the former acting as a membrane transporter on the lysosomal membrane and the latter as a soluble transporter in the lumen. Currently, NPC2 is thought to bind to luminal cholesterol and transfer it to NPC1’s luminal N-terminal domain (NTD), which then transports cholesterol to the lysosomal membrane or for egress (22). NPC2 is sufficient for cholesterol transport between liposomes *in vitro* (23), so its interaction with NPC1 has been proposed to be required in the overcome membrane shielding by luminal glycans in the lysosome (22, 24). In yeast, loss of both Ncr1, the homolog of NPC1, and Npc2 has been observed to reduce the frequency of vacuole microdomains, with the effect greater for the latter (25).

Here, we explore the pathways by which sphingolipids are transported to the vacuole to enable microdomain formation. Using a fluorescent ceramide analogue, we provide evidence that sphingolipid trafficking to the vacuole is mediated by endosomal sorting pathway. We further demonstrate that Npc2 functions as an intravacuolar sphingolipid transporter, acting independently of Ncr1.

## Methods

### Yeast strains and plasmids

The wildtype yeast used in this study is W303a. Using PCR-amplified cassettes, knockouts were generated by homologous recombination via the lithium acetate transformation method. Fluorescent protein markers were expressed either by genetic integration using an integration plasmid or by episomal expression from multicopy plasmids. The gene fragment for yeast codon-optimized human NPC2 was synthesized (Genscript) and inserted into the yeast genome by homologous recombination in replacement of endogenous yeast *NPC2* locus. A list of strains and plasmids can be found in Table S1 and Table S2, respectively.

### Cell growth

To induce vacuole domains, cells were grown as described previously (1). Briefly, a single colony was initially inoculated in nutrient-rich yeast peptone dextrose (YPD). After 18 hours of incubation, cells were diluted into Complete Synthetic Media (CSM) with 2% (w/v) glucose. After an additional 18 hours of incubation, cells were diluted into minimal media with 0.4% (w/v) glucose containing only essential amino acids and nucleotide bases.

### Microscopy and analysis

For fluorescence microscopy, 200 µL of yeast culture was added to wells of an 8-well microscope chamber slide (Nunc Lab-Tek, Thermo Fisher Scientific) pretreated with 200 µL of 2mg/mL concanavalin A (MP Biomedicals) to immobilize the cells. Images were acquired using a Zeiss LSM 880 equipped with a 63x/1.4 Plan-Apochromat oil objective and an Airyscan detector. For BODIPY C12-Ceramide (BODIPY-Cer) imaging, samples were excited using a 488nm Argon laser set at 2.0% power, and images were acquired using ZEN BLACK with 8x averaging and processed using default Airyscan settings. For BODIPY-Cer imaging, samples were excited using a 488nm Argon laser set at 2.0% power and images were acquired with 8x line averaging and processed using the default Airyscan settings. For dual-staining experiments with BODIPY-Cer and FM 4-64 ((*N*-(3-Triethylammoniumpropyl)-4-(6-(4-(Diethylamino) Phenyl) Hexatrienyl) Pyridinium Dibromide), Invitrogen), images were acquired in LSM mode using a 32-channel GaAsP detector. Excitation was performed with a 488 nm Argon laser, with emissions collected from 493nm-589 nm for BODIPY-Cer, and 600-758 nm for FM 4-64.

Vacuole domain frequency was assessed manually by examining the entire Z-stack of widefield images of yeast expressing the vacuolar Ld marker, GFP-Pho8 in ImageJ. Each vacuole was classified as uniform, non-uniform or no domains using the multipoint tool in ImageJ. Uniform vacuoles display evenly distributed vacuole domains, often arranged in hexagonal array; non-uniform vacuoles have irregularly spaced vacuole domains, and vacuole with no domains lack regions excluding GFP-Pho8 at the resolution limit of Airyscan detector. A minimum of 100 vacuoles were counted per biological replicate.

Vacuole domain size was measured using Weka segmentation. For this, Z-stack fields of yeast vacuoles were cropped into individual vacuole micrographs. From each cropped vacuole, a single slice was chosen by visual inspection to ensure the clearest visualization of domains. These slices were compiled, and size analysis was carried out using Trainable Weka Segmentation (26). Vacuole domains were initially classified according to a previously established training set annotated into three categories: background, unlabeled Lo domains, and Ld domain marked by GFP-Pho8 (27). Domains that were not captured by the trained Weka model were manually annotated and incorporated into the training set to improve accuracy. Segmented domains were then quantified using the ImageJ particle analysis tool, followed by final visual inspection to remove false positives arising from mis-segmentation.

### BODIPY C12-Cer staining

BODIPY C12-Ceramide (d18:1/12:0) was purchased from Cayman Chemical (item no. 25997). The stock solution was prepared to 1 mg/mL concentration in Chloroform:Methanol (19:1, v/v) mixture. 50 μL of the stock solution was transferred to a glass tube and dried under a stream of nitrogen gas. To evaporate any remaining solvent, the vial was dried under vacuum for at least 2 hours. The dried BODIPY C12-Ceramide film was resuspended in 200 μL of absolute ethanol, transferred to a microcentrifuge tube and stored protected from light.

For staining, 1 mL of yeast culture (grown as described above), was collected into a 1.5mL sterile microcentrifuge tube and pelleted at 3,000 ×*g* for 1 minute. The supernatant was transferred to a separate tube, combined with 10 µL of 3.4 mg/mL BSA in 0.4% (w/v) glucose minimal media and vortexed. Then, 20 μL of BODIPY C12-ceramide solution was added and vortexed again to generate a BSA-BODIPY C12-ceramide complex. This complex was added back to the cells, and incubated for 1 hr. In parallel, the supernatant of remaining culture (4mL) was saved to resuspend the cells after washing in order to minimize metabolic change. After one hour, cells were washed three times with sterile MilliQ water and resuspended in 1mL of the saved supernatant of the remaining culture, before imaging.

### Npc2 expression and purification

To minimize the protease activity, a protease deficient strain, *pep4Δprb1*Δ was used for expression. The native yeast NPC2 gene was cloned into an expression construct under the control of the *GAL1* promoter with a C-terminal purification tag containing a thrombin cleavage site followed by a deca-histidine tag as described previously (22). For expression, Npc2 expressing yeast grown in CSM with 2% (w/v) glucose were diluted into 650 mL of 0.4% glucose (w/v) minimal media under selection to final OD_600_ of 0.2, incubated for approximately 8 hours until the culture reached OD_600_ of 0.8-1.0 per mL. Npc2 expression was induced by adding galactose to a final concentration of 2% (w/v) followed by incubation for 22 hours. Cells were harvested by centrifugation at 3000 ×*g* for 30 min at 4 °C, washed with ice-cold water for 3 times, and pellets were stored at −80 °C.

Frozen cell pellets were thawed and resuspended in approximately 75 mL lysis buffer (600 mM NaCl, 100 mM Tris-HCl pH 7.5). 3 mL of 25x complete EDTA-free protease inhibitor (Roche) solution in water was added to the suspension. The cells were lysed using EmulsiFlex C3 (Avestin), at pressure of 1500 bar. 25 mL of the lysis buffer was used to rinse out any remaining cells in the Emulsiflex system and combined with the cell lysate. 1 mL of additional protease inhibitor was added to the lysate. The lysate was centrifuged at 39,000 rpm for 30 minutes at 4 °C using a 45Ti rotor (Beckman) to remove unlysed cells and cell debris. The supernatant was collected and syringe-filtered through 0.2 μm pore to further remove debris before running through a Ni-NTA column (Roche) on a FPLC system (Akta Pure 25). A W10 buffer (500 mM NaCl, 50 mM Tris pH 7.5, 10% glycerol, 10 mM imidazole) was used to equilibrate the column. After the lysates were loaded on to the Ni-NTA column, 20 CV of the W10 buffer was used to wash the column. The proteins were eluted with 500 mM NaCl, 50 mM Tris pH 7.5, 10% glycerol, 500 mM imidazole buffer, and fractions were analyzed on SDS-PAGE gel by staining with instant blue (Abcam). Fractions containing the protein were pooled and concentrated on a protein concentrator (Amicon Ultra 15 Centrifugal filter 3kDa MWCO, Millipore) to 2mL. The sample was loaded on a Sephacryl S200 HR column equilibrated in a gel filtration buffer (200 mM NaCl, 20 mM Tris pH 7.5). The eluate absorbance was monitored at 280 nm, and fractions were collected in 2.0 mL volumes. Fractions corresponding to the protein peak were collected and analyzed by SDS-PAGE gel. Those containing proteins were pooled and concentrated to ∼0.1 mg/mL to use for lipid transport assay. Protein samples were analyzed for purity and mass identification by liquid chromatography with an Agilent 6230 time-of-flight mass spectrometer (TOFMS) with JetStream electrospray ionization source (ESI). Glycerol was added to 10% (v/v) to the remaining protein sample, then frozen in liquid nitrogen for long-term storage.

### Liposome preparation

POPG (1-palmitoyl-2-oleoyl-sn-glycero-3-(phospho-rac-(1-glycerol))), DOPC (1,2-dioleoyl-sn-glycero-3-phosphocholine), Dansyl-PE (18:1 1,2-dioleoyl-sn-glycero-3-phosphoethanolamine-N-(5-dimethylamino-1-naphthalenesulfonyl), ammonium salt), dehydroergosterol (DHE), and Rhod-PE (18:1 1,2-dioleoyl-sn-glycero-3-phosphoethanolamine-N-(lissamine rhodamine B sulfonyl), ammonium salt) were purchased from Avanti Polar Lipids. C12 NBD-Dihydroceramide (DHCer) and C12 NBD-IPC were synthesized from precursors commercially obtained. The stock solutions of each lipid were prepared by dissolving the lipid in chloroform, concentrations ranging from 1 mg/mL to 25 mg/mL. Liposomes were prepared by drying 4 µmol of total lipid in a clean glass tube under a stream of nitrogen gas, followed by at least two hours under vacuum to remove residual solvent. The lipid film was hydrated in 1 mL of buffer CN (20 mM sodium citrate, 150 mM NaCl, pH 5.0) to a final lipid concentration of 4 mM, and vortexed vigorously for 1 minute. The vesicles were agitated for at least 1 hour. The vesicles were then freeze-thawed 5 times and extruded 21 times through a 200 nm polycarbonate membrane using 250 μL mini extruder kit (Avanti Polar Lipids) to produce unilamellar vesicles.

### NBD-DHCer and NBD-IPC synthesis

C12 NBD-DHCer was synthesized via HATU coupling as previously described (28). NBD-IPC was synthesized similarly, using 18:1 sphingosyl-PI (D-erythro-sphingosyl phosphoinositol) (Avanti Polar Lipids) and NBD-dodecanoic acid (C12-NBD) (Avantor Sciences). O-(7-azabenzotriazol-1-yl)-1,1,3,3-tetramethyluronium hexafluorophosphoate (HATU) and *N,N*- diisopropylethylamine (DIPEA) were obtained from Sigma Aldrich. 1.33 mg of C12-NBD (3.51 µmol, 1.0 equiv), 1.47 mg of HATU (3.86 µmol, 1.1 equiv) and 1.83 µL of DIPEA (10.53 µmol, 3.0 equiv) was added into a reaction vial, dissolved in approximately 200 µL of anhydrous dimethylformamide (DMF) and stirred for at least 5 minutes under nitrogen gas. 2 mg of sphingosyl-PI (3.51 µmol, 1.0 equiv) was added to the reaction vial and DMF was added to the total of 500 µL. The reaction vial was flushed with nitrogen gas and the reaction was stirred overnight at room temperature in the dark. DMF was removed by rotary evaporation. The compound was resuspended in methanol and filtered through a 0.2 µm syringe driven PTFE filter before purification by reverse-phase HPLC purification by reverse-phase HPLC on an Agilent 1260 Infinity II LC system fitted with a C18 semipreparative column. Elution was performed using a gradient with Phase A (H₂O containing 0.1% TFA) and Phase B (acetonitrile containing 0.1% TFA). The HPLC program consisted volume/volume ratios of 50:50 (A:B) for 2 min, a linear gradient to 25:75 over 8 min, followed by a gradient to 1:99 over 4 min. Compound identity and purity was confirmed by high resolution LCMS (calculated for C_42_H_73_N_5_O_14_P [M+H]^+^, *m/z* 902.4886, found 902.5119, Thermo Scientific Q-Exactive Orbitrap) and 1H-NMR Bruker Avance 600 NMR at UCSD Biomolecular NMR facility. Yellow oil; 1.26 mg (1.40 µmol, 39.9% yield). Lower yield attribute to incomplete fraction collection during early elution. ^1^H-NMR: δ 0.87 ppm (t, 3H, terminal CH₃), 1.15 ppm (br m, 46H, aliphatic (CH₂)ₙ), 3.31–5.41 ppm (m, 12H, sugar/linker protons), 6.98–7.56 ppm (m, 2H, aromatic/NBD).

### Lipid transport assay

Transport of DHE was performed as described previously (29) using 1 μM concentration of purified yeast Npc2. The donor liposome (L_D1_) was composed of 30 mol % POPG, 62.5 mol % DOPC, 5 mol % DHE and 2.5 mol % Dansyl-PE; the acceptor liposome (L_A1_) was composed of 30 mol % POPG and 70 mol % DOPC. To determine baseline fluorescence (F_0_) for fluorescence normalization, the L_D-No DHE_ liposomes were prepared with 30 mol % POPG, 67.5 mol % DOPC and 2.5 mol % Dansyl-PE. In each experiment, 200 µM L_A1_ was added to the cuvette filled with buffer CN with a small magnetic stir bar. After 1 minute, 200 µM L_D1_ was added. After 4 minutes, 1 µM Npc2 was added. For control experiments, equal volume of buffer was added instead of protein. The fluorescence reading was monitored over a 30-minute interval with continuous stirring. The fluorescence values were normalized using the following equation: Normalized fluorescence = 1 - (F - F_0_) / (F_max_ - F_0_), where F is the fluorescence intensity at each time point, F_0_ corresponds to the average fluorescence reading over 5 minutes after 9 minutes of incubation L_D- No DHE_ and L_A1_, and F_max_ is the maximum fluorescence determined by averaging the fluorescence intensity from the 6-second window immediately prior to the addition of Npc2. The concentration of DHE transported was calculated by multiplying normalized fluorescence by 10, reflecting the full 10 µM DHE pool accessible for transfer due to rapid sterol flip-flop between bilayer leaflets. The fluorescence readings were taken on a Cary Eclipse fluorometer (Agilent) in kinetics mode. The excitation wavelength was set to 310 nm, and the emission wavelength was set to 525 nm to observe Forster Resonance Energy Transfer (FRET) between DHE and Dansyl-PE. The sample holder was temperature controlled at 30 °C.

For NBD-DHCer and NBD-IPC transport assay, the donor liposome (L_D2_) was prepared with 30 mol % POPG, 65 mol % DOPC, 5 mol % NBD-DHCer or NBD-IPC, and the acceptor liposome (L_A2_) was prepared with 30 mol % POPG and 68 mol % DOPC, and 2 mol % Rhod-PE. In order to normalize the fluorescence, donor liposome at equilibrium (L_D-eq_) consisting of 30 mol % POPG, 67.5 mol % DOPC and 2.5 mol % NBD-DHCer, and acceptor liposome at equilibrium (L_A-eq_) vesicles consisting of 30 mol % POPG, 65.5 mol % DOPC, 2.5 mol % NBD-DHCer and 2 mol % Rhod-PE were also prepared. In each experiment, 200 µM L_D2_ liposomes were added to the cuvette, followed by addition of 200 µM L_A2_ liposomes after 1 minute. After 4 minutes, 1 µM Npc2 was added to the cuvette, and the fluorescence readings were taken continuously over 30 minutes with magnetic stirring. The fluorescence reading was normalized using the formula: Normalized fluorescence = 1 - (F - F_eq_) / (F_max_ - F_eq_), where F_eq_ is average fluorescence signal of two population of liposomes L_A-eq_ and L_D-eq_ and F_max_ is the average fluorescence of L_A_ and L_D_ over 6 seconds right before addition of Npc2. The concentration of NBD-IPC transported to the acceptor liposome was calculated by multiplying the normalized fluorescence by 5 which corresponds to the concentration (µM) of NBD-lipid accessible on the outer leaflet of liposomes (28). For NBD- DHCer, transport was expressed as percent of total lipid transferred rather than concentration due to the high melting temperature of NBD-DHCer, which caused loss of fluorescent lipids at the filter support during extrusion and led to unreliable values after normalization with equilibrium liposomes. The excitation wavelength was set to 473 nm, and the emission wavelength was set to 538 nm to monitor the fluorescence.

### Vacuole isolation

Vacuoles were isolated from yeast in the early stationary phase using the method described previously (30). Yeast cells grown in 650 mL of 0.4% (w/v) glucose minimal media were harvested at 3,000 ×*g* and used fresh for vacuole isolation. After incubating in 100 mM Tris-HCl, pH 9.5 with DTT, the cells were washed multiple times. The cells were then treated with 9 mg zymolyase 20T per wet weight cell for 1 hour at 30 °C to remove the cell wall, mechanically lysed using Dounce homogenizer to liberate the vacuole. Through a series of density gradient centrifugation using Ficoll 400, the vacuoles were isolated from the top layer of the gradient. To further remove other organelle contaminants, the vacuoles were subjected to a gravity centrifugation and were collected at the very bottom of the tube. The isolated vacuoles were used for lipidomics analysis.

### Lipidomics

Mass spectrometry-based global lipid analysis was performed by Lipotype GmbH (Dresden, Germany) as described (31, 32). Lipids were extracted using a two-step chloroform/methanol procedure (31). Samples were spiked with internal lipid standard mixture containing: CDP-DAG 17:0/18:1, cardiolipin 14:0/14:0/14:0/14:0 (CL), ceramide 18:1;2/17:0 (Cer), diacylglycerol 17:0/17:0 (DAG), lyso-phosphatidate 17:0 (LPA), lyso-phosphatidyl-choline 12:0 (LPC), lyso-phosphatidylethanolamine 17:1 (LPE), lyso-phosphatidylinositol 17:1 (LPI), lyso-phosphatidylserine 17:1 (LPS), phosphatidate 17:0/14:1 (PA), phosphatidylcholine 17:0/14:1 (PC), phosphatidylethanolamine 17:0/14:1 (PE), phosphatidylglycerol 17:0/14:1 (PG), phosphatidylinositol 17:0/14:1 (PI), phosphatidylserine 17:0/14:1 (PS), ergosterol ester 13:0 (EE), triacylglycerol 17:0/17:0/17:0 (TAG), stigmastatrienol, inositolphosphorylceramide 44:0;2 (IPC), mannosyl-inositolphosphorylceramide 44:0;2 (MIPC) and mannosyl-di-(inositolphosphoryl)ceramide 44:0;2 (M(IP)_2_C). After extraction, the organic phase was transferred to an infusion plate and dried in a speed vacuum concentrator. 1^st^ step dry extract was re-suspended in 7.5 mM ammonium acetate in chloroform/methanol/propanol (1:2:4, V:V:V) and 2^nd^ step dry extract in 33% ethanol solution of methylamine in chloroform/methanol (0.003:5:1; V:V:V). All liquid handling steps were performed using Hamilton Robotics STARlet robotic platform with the Anti Droplet Control feature for organic solvents pipetting.

Samples were analyzed by direct infusion on a QExactive mass spectrometer (Thermo Scientific) equipped with a TriVersa NanoMate ion source (Advion Biosciences). Samples were analyzed in both positive and negative ion modes with a resolution of R_m/z=200_=280000 for MS and R_m/z=200_=17500 for MS-MS experiments, in a single acquisition. MS-MS was triggered by an inclusion list encompassing corresponding MS mass ranges scanned in 1 Da increments (33). Both MS and MS-MS data were combined to monitor EE, DAG and TAG ions as ammonium adducts; PC as an acetate adduct; and CL, PA, PE, PG, PI and PS as deprotonated anions. MS only was used to monitor CDP-DAG, LPA, LPE, LPI, LPS, IPC, MIPC, M(IP)_2_C as deprotonated anions; Cer and LPC as acetate adducts and ergosterol as protonated ion of an acetylated derivative (34). Data were analyzed with in-house developed lipid identification software based on LipidXplorer (35, 36). Data post-processing and normalization were performed using an in-house developed data management system. Only lipid identifications with a signal-to-noise ratio >5, and a signal intensity 5-fold higher than in corresponding blank samples were considered for further data analysis.

## Results

### Ceramide is routed to the vacuole via endosomal sorting pathways

The trafficking pathways for membrane proteins from the Golgi to plasma membrane to endosomes and vacuole require Vps proteins. Previously, *vps* mutants have been categorized based on their vacuole morphology based on immunofluorescence microscopy with antibodies against vacuole protein, ALP (37). Type A mutants contain WT-like morphologies, which include single large vacuoles as cells enter the stationary stage. In contrast, *vps4*Δ cells which were previously shown to have reduced domain (2), are classified as a Type E mutant and show extensive vacuole fragmentation. We have observed that fragmented vacuoles uniformly do not exhibit domains (27). However, vacuole fusion alone is not sufficient to drive domain formation, as *AUR1* knockdown cells with reduced sphingolipid synthesis display neither domains nor increased fragmentation (1).

To identify specific mechanisms of sphingolipid sorting to the vacuole, we first screened the distribution of GFP-conjugated ALP (GFP-Pho8), a vacuole Ld domain marker, in eight previously identified Type A *vps* mutants. While *vps13*Δ, *vps29*Δ, *vps30*Δ cells had clear differences in vacuole domain formation (Figure 1A), vacuoles from *vps8*Δ, *vps10*Δ and *vps43*Δ cells appeared similar to that of WT (Figure S1). The observation of normal vacuole domains *vps10*Δ, which lack the binding receptor required for CPY pathway, suggested that CPY pathway is not a primary pathway for trafficking of domain-promoting lipids to the vacuole. Loss of Vps13, a bridge-like lipid transport protein that acts at multiple organelle contact sites (38), led to more fragmented vacuoles, which showed no phase separation. We thus focused further analysis on *vps29*Δ and *vps30*Δ. Vps29 is a retromer subunit involved in endosome to Golgi retrograde transport (39). While *vps29*Δ cells showed a high frequency of uniform vacuole domains (Figure 1B), they were smaller in size (Figure 1C) and often characterized by small puncta of GFP-Pho8 near the vacuole. The latter was also observed in *vps35*Δ, another retromer subunit (Figure S1). Vps30, also known as Atg6, is a subunit of phosphatidylinositol 3-kinase complexes I and II that are essential for autophagy and vps pathways, respectively (40). In *vps30*Δ cells, most vacuoles had non-uniform vacuole domain formation (Figure 1B), which we had previously observed in mutants with altered sphingolipid headgroup distributions (1).

**Figure 1.**
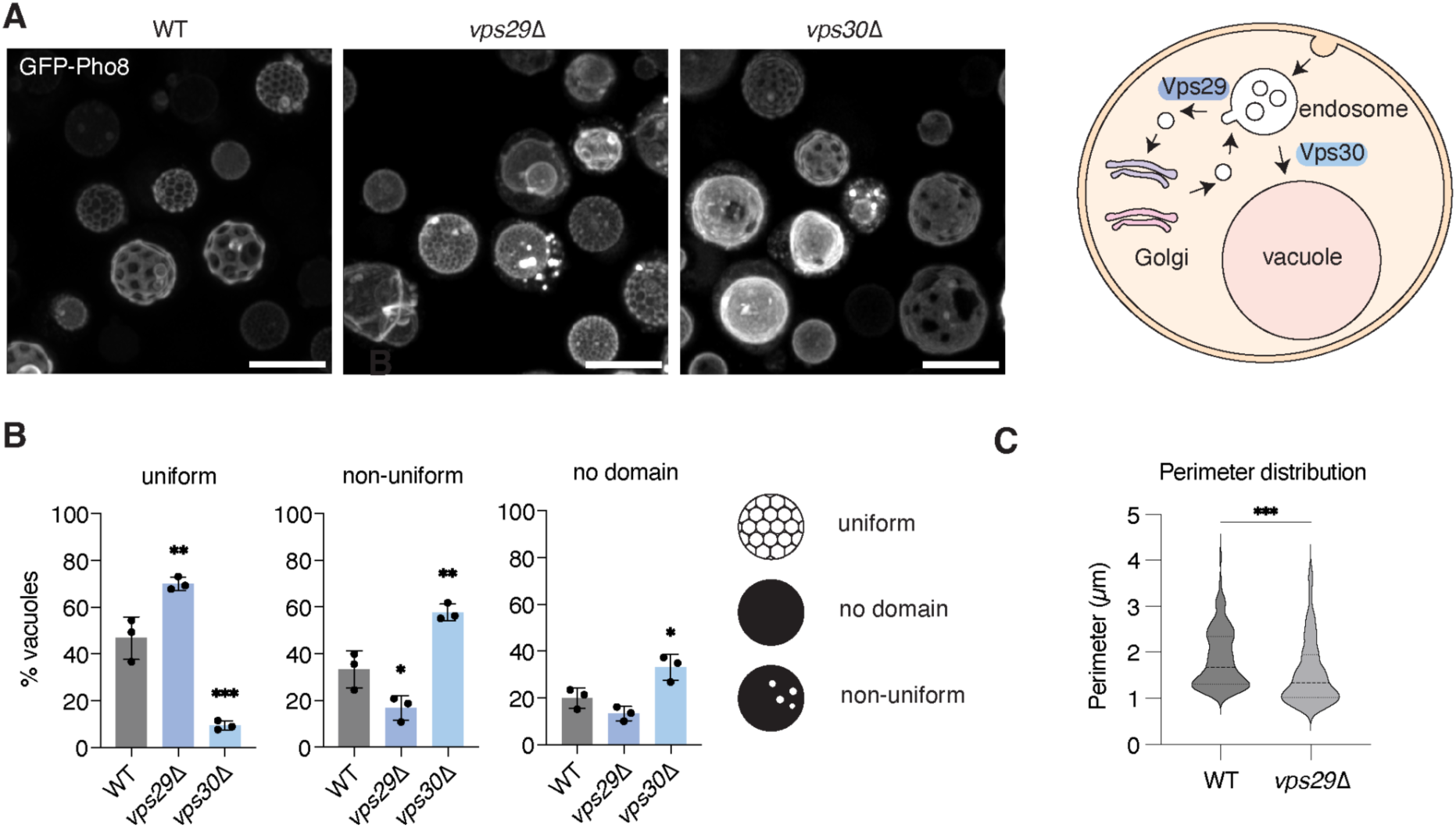
Two Type A mutants (*vps29*Δ and *vps30*Δ) show distinct vacuole morphology compared to the WT cells. **A.** Representative field view of vacuoles visualized by expression of Ld marker, GFP-Pho8. Scale bar, 5 µm. **B.** Quantification of vacuole domain types. Cells lacking Vps29 display more uniform vacuole domains, while those lacking Vps30 feature more non-uniform vacuole domains (n = 3 individual cultures, N > 130 cells for each). Significance against WT was assessed by one-way ANOVA with Dunnett’s post-hoc test; *p < 0.05; **p < 0.01; ***p < 0.001. **C.** Cells lacking Vps29 show smaller vacuole domains, as measured by their perimeter lengths, than WT cells. Significance was assessed by unpaired two-tailed t-test against the WT; ***p < 0.001.

To test whether *vps* mutants altered sphingolipid trafficking, we employed live cell imaging with a fluorescent sphingolipid analogue. Although ceramides are less abundant than complex sphingolipids like IPC in the vacuole membrane, they also become enriched during microdomain formation, and short chain ceramides can be added to cells externally to label endomembranes. In contrast to the more commonly used ceramide analogue NBD-C6-Ceramide, we found that BODIPY-C12-Ceramide (BODIPY-Cer, Figure S2A) is incorporated into live yeast cells and stains the vacuole membrane – including its Lo domains – after extended incubations (Figure S2B). Analysis by TLC confirmed that BODIPY-Cer is not modified nor degraded under our experimental conditions (Figure S2C), so it can be used as a read out of ceramide trafficking, rather than metabolism. We therefore used this probe to visualize changes in ceramide distributions that could be relevant for broader sphingolipid trafficking.

We observed changes to BODIPY-Cer distributions in cells lacking both Vps29 and Vps30, the Type A mutants showing altered domain formation capacity. In *vps29*Δ cells, there was an abundance of smaller BODIPY-Cer puncta near the periphery of the cell (Figure 2A). Similarly, *vps30*Δ cells showed more BODIPY-Cer puncta near the plasma membrane, but also lining the vacuole. To test the identity of these puncta, we co-stained cells for short time periods (1 hour) with the dye FM 4-64, which is internalized through endosomes and transported to the vacuole (41). FM 4-64 co-localized with BODIPY-Cer puncta peripheral to the vacuole, as well as BODIPY-Cer signal near the PM in *vps29*Δ and *vps30*Δ cells. In contrast, the autophagosome marker Atg8-RFP did not show co-localization with BODIPY-Cer (Figure S3A). Lipid droplets, marked by Erg6-BFP, showed partial co-localization with some of the larger BODIPY-Cer aggregates, likely due to the hydrophobicity of the stain (Figure S3A). However, the smaller punctae that lined the vacuole membrane did not show co-localization in Erg6-BFP (Figure S4A). These experiments suggested that externally added BODIPY-Cer enters the vacuole through endosomal sorting, which is impeded in *vps* mutants.

**Figure 2.**
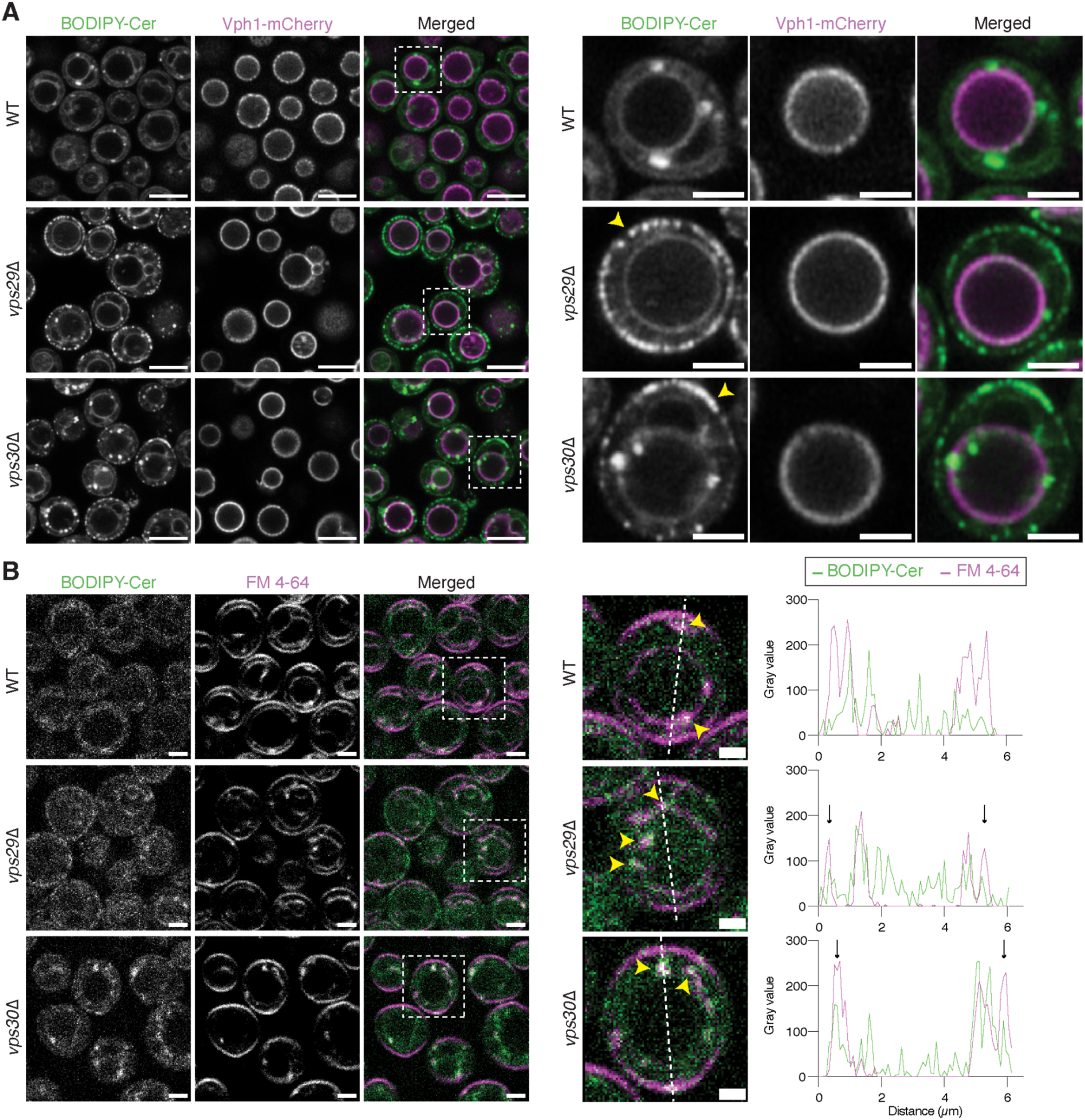
Alterations in sphingolipid trafficking imaged with BODIPY-Cer. **A.** Images of VPS mutants expressing a vacuole marker, Vph1-mCherry stained with BODIPY-Cer taken 1 hour after staining. Scale bar, 5 µm. Both *vps29*Δ and *vps30*Δ cells show punta around the plasma membrane, which is not observed in WT cells. **B.** Cells were co-stained with BODIPY-Cer and FM 4-64. Insets highlight colocalization of intracellular BODIPY-Cer puncta with FM 4-64 (yellow arrows). The line profile on the right shows colocalization of BODIPY-Cer and FM 4-64 in bright punctae, indicating endosomal trafficking of exogenously added sphingolipids. In WT cells, colocalization is observed at endosomes near the vacuole, while *vps29*Δ and *vps30*Δ also show accumulation peripheral to the PM (black arrows).

### Npc2 mediates sphingolipid egress from the vacuole lumen

We next asked how sphingolipids are incorporated into the vacuole membrane from endosomal cargoes. We were motivated by previous observations that expansion of vacuole microdomains was dependent on the yeast NPC proteins Ncr1 and Npc2 (25). This effect could be due to defects in vacuole incorporation of ergosterol, which also increases in abundance upon microdomain formation but to a lesser extent than sphingolipids. However, yeast Npc2 has been shown to bind a range of lipids in addition to sterols (42), including phosphatidylinositol which bears structural similarities to the sphingolipid IPC (43). We thus hypothesized that the NPC system could play a broader role in vacuole sphingolipid trafficking. Under our experimental conditions, cells lacking Npc2 (*npc2*Δ) showed a 38% reduction in domain frequency in GFP-Pho8 expressing cells (Figure 3A), while those lacking Ncr1 (*ncr1*Δ) showed no change. The dominant effect of Npc2 on vacuole domains was consistent with previous work (25).

**Figure 3.**
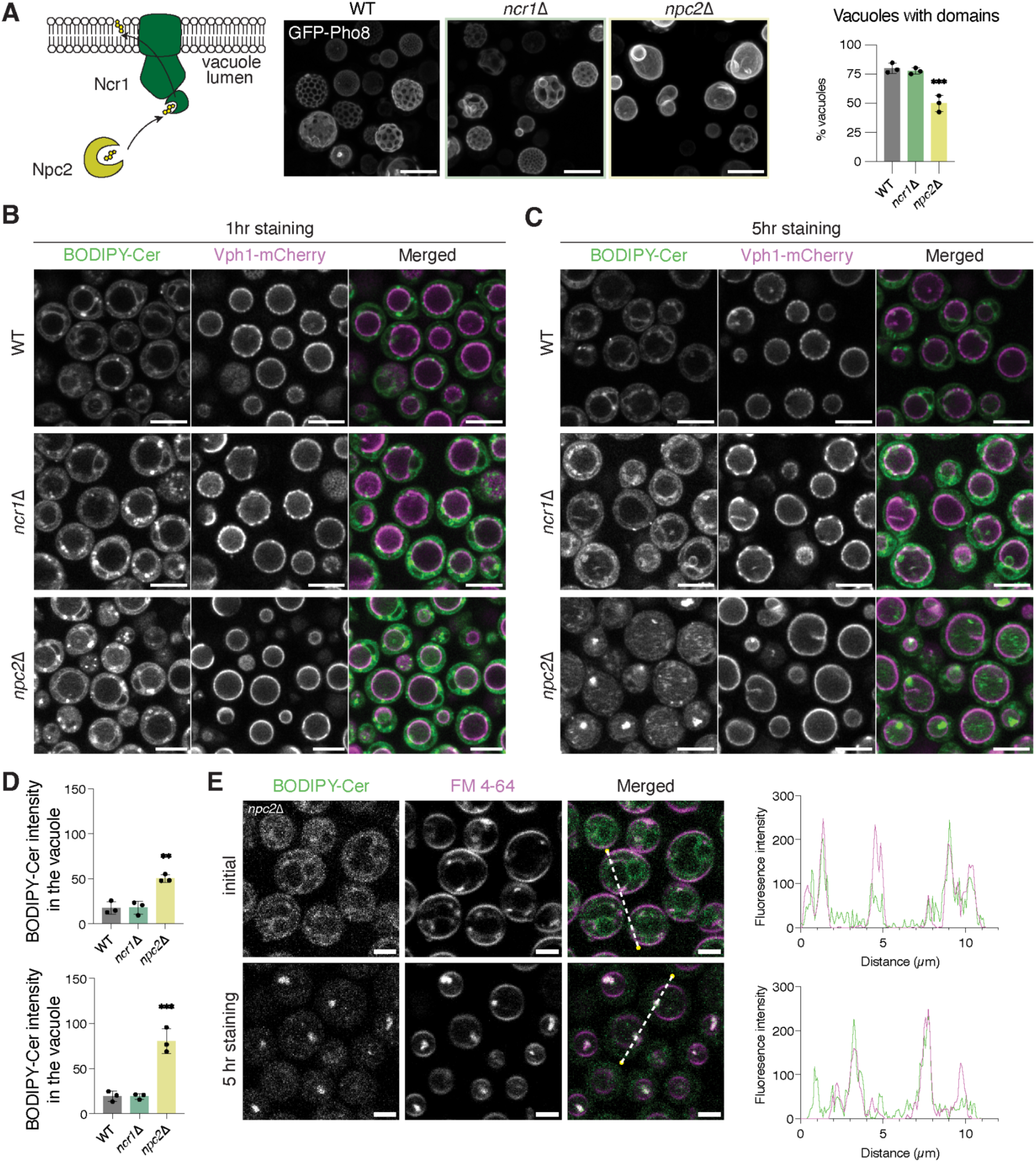
A role for Npc2 in vacuole membrane incorporation of intraluminal sphingolipids. **A.** Analysis of vacuole microdomain formation in mutants of two NPC proteins Ncr1 and Npc2. Only *npc2*Δ shows a loss of vacuole domains. Significance against WT was assessed by one-way ANOVA with Dunnett’s post-hoc test; ***p < 0.001, n = 3 independent cultures. **B.** BODIPY-Cer distributions in ncr*1*Δ and *npc2*Δ cells after 1 hr of staining. Both look similar to WT, with the appearance of internalized BODIPY-Cer signals in *npc2*Δ. Scale bar, 5µm. **C.** After 5 hours of staining, *npc2*Δ cells show widespread intraluminal BODIPY-Cer puncta that is not observed in WT or *ncr1*Δ. Scale bar, 5 µm. **D.** Quantification of internalized BODIPY-Cer signal in the vacuole of *ncr1*Δ and *npc2*Δ cells, with the latter having higher intensity compared to WT that increases over time. Significance against WT was assessed one-way ANOVA with Dunnett’s post-hoc test; **p < 0.01; ***p < 0.001. n = 3 independent cultures. **E.** Line profiles across *npc2*Δ vacuoles showing that extraluminal (top, 1 hr) and intraluminal (bottom, 5 hr) BODIPY-Cer puncta colocalize with the endosomal marker FM 4-64. Scale bar, 2 µm.

We stained NPC mutant cells with BODIPY-Cer to identify any possible alterations in sphingolipid trafficking. While the pattern of BODIPY-Cer in *ncr1*Δ resembled that of WT cells, *npc2*Δ cells showed a striking accumulation of BODIPY-Cer signals as aggregates appearing within vacuole lumens (Figure 3B). This intraluminal signal increased as BODIPY-Cer incubations were extended from 1 to 5 hours (Figure 3C). In contrast, *ncr1*Δ cells showed no increase in internalized BODIPY-Cer signal relative to WT. The BODIPY-Cer punctae in *npc2*Δ cells co-localized with the endosomal marker FM 4-64 (Figure 3E), but not Atg8-RFP (Figure S3B) or Erg6-BFP (Figure S4B). Thus, cells lacking Npc2 accumulated BODIPY-Cer within the vacuole that is likely to be endosomal in origin.

To confirm that Npc2, independently plays a role in sphingolipid sorting into the vacuole membrane, we purified early stationary stage vacuoles from both *npc2*Δ and *ncr1*Δ cells using a density separation methodology we previously developed (30). Subsequent lipidomic analysis revealed that early stationary stage vacuoles from *npc2*Δ cells contained lower amounts of all sphingolipids, including ceramides, and ergosterol compared to those from WT cells (Figure 4B, 4C). In contrast, *ncr1*Δ vacuoles showed a redistribution of sphingolipid species – an increase in MIPC at the expense of M(IP)_2_C (Figure 4C) and changes to their length and hydroxylations (Figure S5A) – but no overall change in their total levels. There were also subtle changes in the glycerophospholipid content of the vacuole, with accumulation of phosphatidylinositol (PI) and decrease of phosphatidylcholine (PC) in both mutants, as well as a slight imbalance in phosphatidylethanolamine (PE) and an accumulation of PS in *ncr1*Δ (Figure S5B). Vacuole lipidomes from all strains contained only trace amounts of phosphatidic acid (PA), a characteristic feature of purified vacuole lipidomes (44, 45).

**Figure 4.**
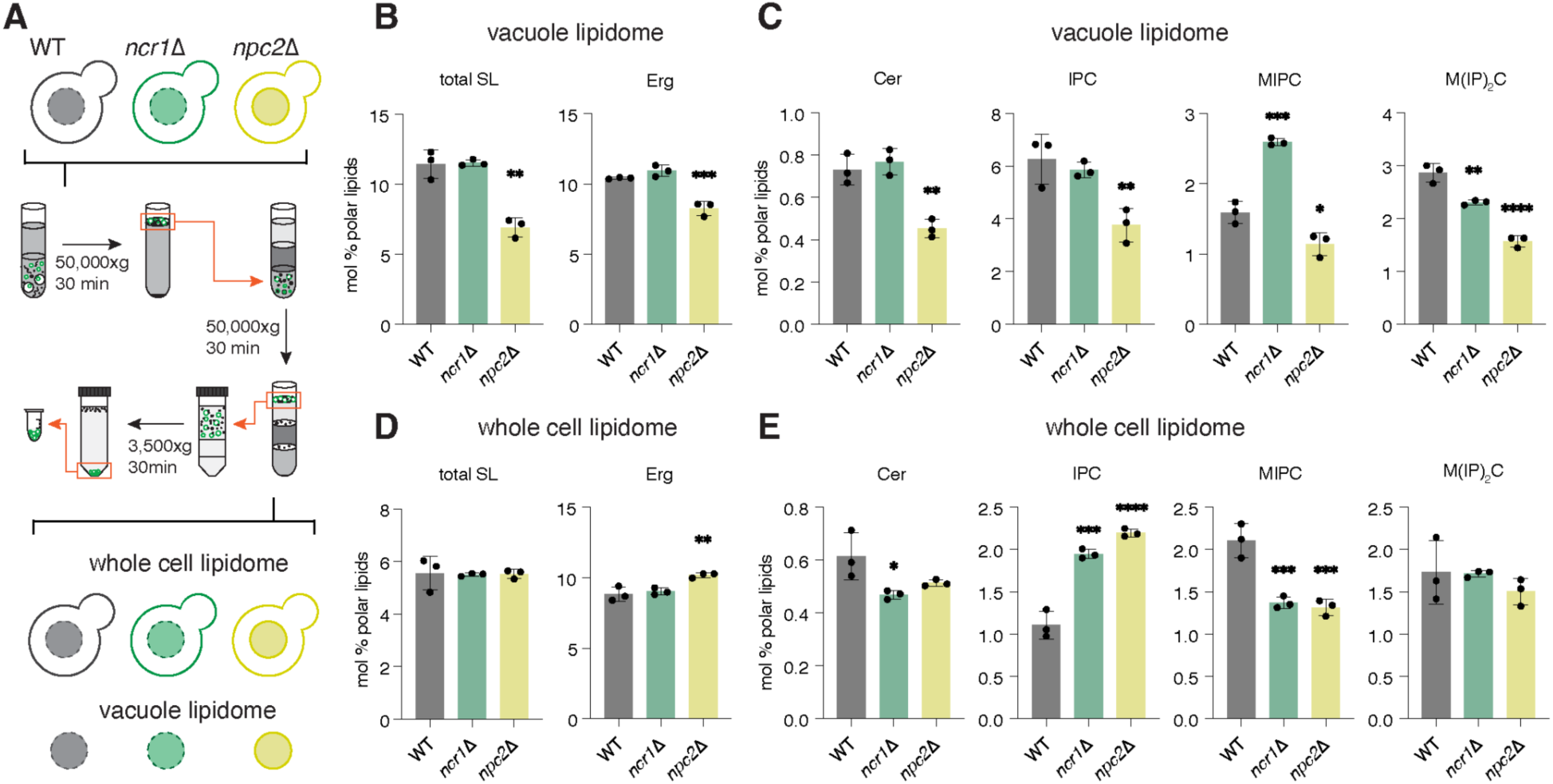
Loss of Npc2 decreases incorporation of sphingolipids in phase-separated vacuole membranes. **A.** Schematic for vacuole isolation from WT and NPC mutant vacuoles. Vacuole and whole cell lipidomes are compared from parallel analyses derived from early stationary stage cells, 24 hours after dilution into 0.4% glucose minimal medium. **B.** Vacuoles from *npc2*Δ cells show reduced levels of sphingolipids, as well as ergosterol, components of Lo membrane domains. **C.** Analysis of individual sphingolipid classes shows reductions in each. **D.** Whole cell lipidomes do not show decreases in total sphingolipid or ergosterol levels but do show changes in the distribution of individual sphingolipid species (**E**). For all panels, Significance against WT was assessed one-way ANOVA with Dunnett’s post-hoc test; *p < 0.05; **p < 0.01; ***p < 0.001, ****p < 0.0001, n = 3 vacuole purification preparations from independently grown cultures or individually grown cell pellets from the same strain grown under identical growth conditions.

In parallel analyses of the corresponding whole cell lipidomes, neither *ncr1*Δ nor *npc2*Δ samples showed altered total sphingolipid levels (Figure 4D). Both mutants showed an increase in IPC levels, with a corresponding decrease in MIPC and ceramide, while M(IP)_2_C remained stable (Figure 4E). Both mutants also showed changes in sphingolipid length and hydroxylation (Figure S5C) and subtle changes in the whole cell phospholipid content (Figure S5D): an increase PA and decrease of phosphatidylethanolamine. Overall, these data suggest that Npc2, but not necessarily its canonical receptor Ncr1, plays a role in vacuole sorting of raft-forming lipids, i.e. ergosterol and sphingolipids, at the onset of membrane phase separation. In contrast, loss of either component has overall effects on sphingolipid metabolism.

### Npc2 acts an independent sphingolipid transporter in liposomes

Given that a range of vacuole sphingolipid phenotypes of *npc2*Δ – microdomain formation, intraluminal BODIPY-Cer accumulation, and membrane lipid composition – are not observed in *ncr1*Δ, we asked if Npc2 acts as a self-sufficient transporter of yeast sphingolipids. To address this possibility, we purified Npc2 from yeast cells for use in liposome-based LTP experiments (Figure S6). In such assays, the transfer of lipids between two liposome populations after protein addition is monitored by FRET. For this, a fluorescent lipid ligand analogue is used as a FRET donor and a second lipid as a FRET acceptor. Regardless of lipid ligand, we observed that yeast Npc2 required an acidic buffer (sodium citrate, pH 5.0) for robust activity, as previously noted for human NPC2 (46), and anionic lipids (30 mol % of phosphatidylglycerol (PG) in phosphatidylcholine (PC) background. Although PG is not abundant in the vacuole, its abundance in the assay matched that of total anionic lipids in the vacuole, including the three complex sphingolipids, PI, and PS, based on our lipidomics data.

We tested the activity of yeast Npc2 to transfer three fluorescent lipids between PC/PG liposomes (Figure 5). For each, 1 µM of protein was added to initiate transfer. The first, dihydroergosterol (DHE), is a fluorescent analogue of ergosterol, containing one extra double bond in the C-ring of the sterol. DHE was first shown to be a substrate for bovine NPC2 (47), and we confirmed its transfer between liposomes that increased upon addition of 1 µM Npc2. The second, NBD-C12-dihydroceramide (NBD-DHCer), is an analogue of ceramide. It showed a higher, Npc2-dependent initial rate of transfer compared to DHE one. Finally, we synthesized a new NBD-labeled analogue of IPC (NBD-IPC), the sphingolipid most highly enriched during microdomain formation (Figure S7). Npc2 robustly transported NBD-IPC (V_i_ = 1.41 ± 0.03 min^-1^ per protein). For both NBD-DHCer and NBD-IPC, FRET-based quenching between the ligand and lipids in acceptor vesicles was used to ensure that the assay measured transport of the ligand, not just extraction and binding. Thus, Npc2 can directly transfer a range of lipids associated with microdomain formation, including the sphingolipid class most strongly sorted into the vacuole membrane under these conditions.

**Figure 5.**
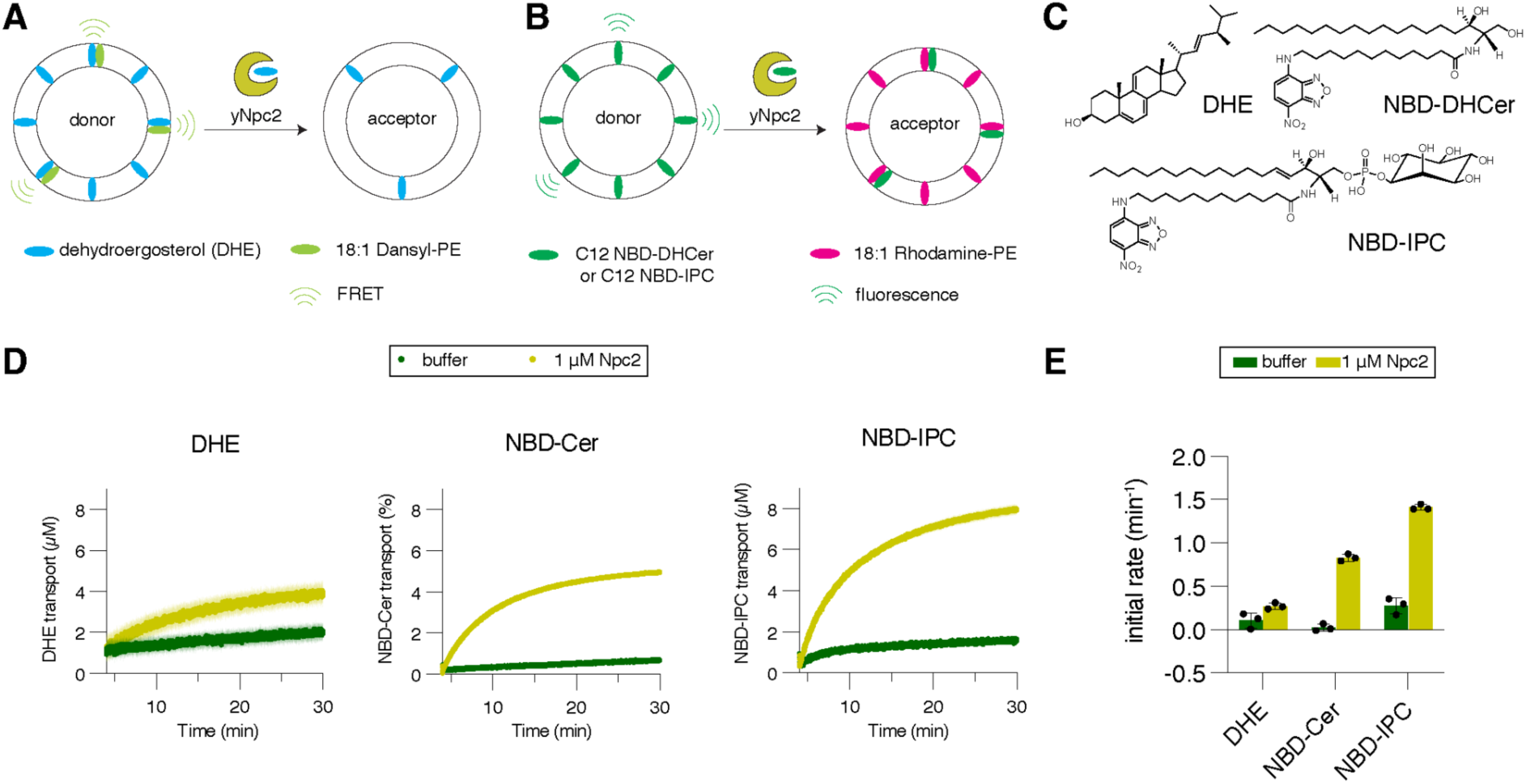
Transport of sphingolipids between liposomes by Npc2. **A.** Schematic for measuring transport of the ergosterol analogue DHE between liposomes based on reduction of the FRET between DHE and Dansyl-PE. Donor vesicles: DOPC/POPG/DHE/Dansyl-PE (62.5/30/5/2.5, mol %). Acceptor vesicles: DOPC/POPG (70/30). **B.** Schematic for measuring transport of sphingolipid analogues NBD-DHCer and NBD-IPC between liposomes based on measurement of quenching of NBD fluorescence with the acceptor Rhodamine-PE. Donor vesicles: DOPC/POPG/NBD-DHCer or NBD-IPC (65/30/5). Acceptor vesicles: DOPC/POPG/Rhod-PE (67.5/30/2.5). **C.** Structures of the analogues used in the assays. **D.** Normalized transport curves for the three substrates. For each, time courses are shown starting upon addition of 1 µM Npc2 or buffer. Plotted are the values for three individual experiments with each substrate. **E.** Initial rates of transport for the three substrates during the first minute after protein or buffer addition.

The capacity for Npc2 to traffic sphingolipids could be a specialized feature of the yeast protein. Human NPC2 (hNPC2) has a small binding cavity that could expand only enough to accommodate smaller lipids like cholesterol (47). In contrast, yeast Npc2 features a larger cavity (Figure 6A) (22) and has recently been shown to bind to polar lipids, like phospholipids, in addition to sterols (43). Consistent with an expanded role for yeast Npc2 in vacuole trafficking, we found that heterologous expression of hNPC2 does not rescue the loss of microdomains in *npc2*Δ yeast (Figure 6B). In contrast, yeast Npc2 had been previously shown to phenotypically rescue loss of cholesterol transport in fibroblasts expressing mutant hNPC2 (48).

**Figure 6.**
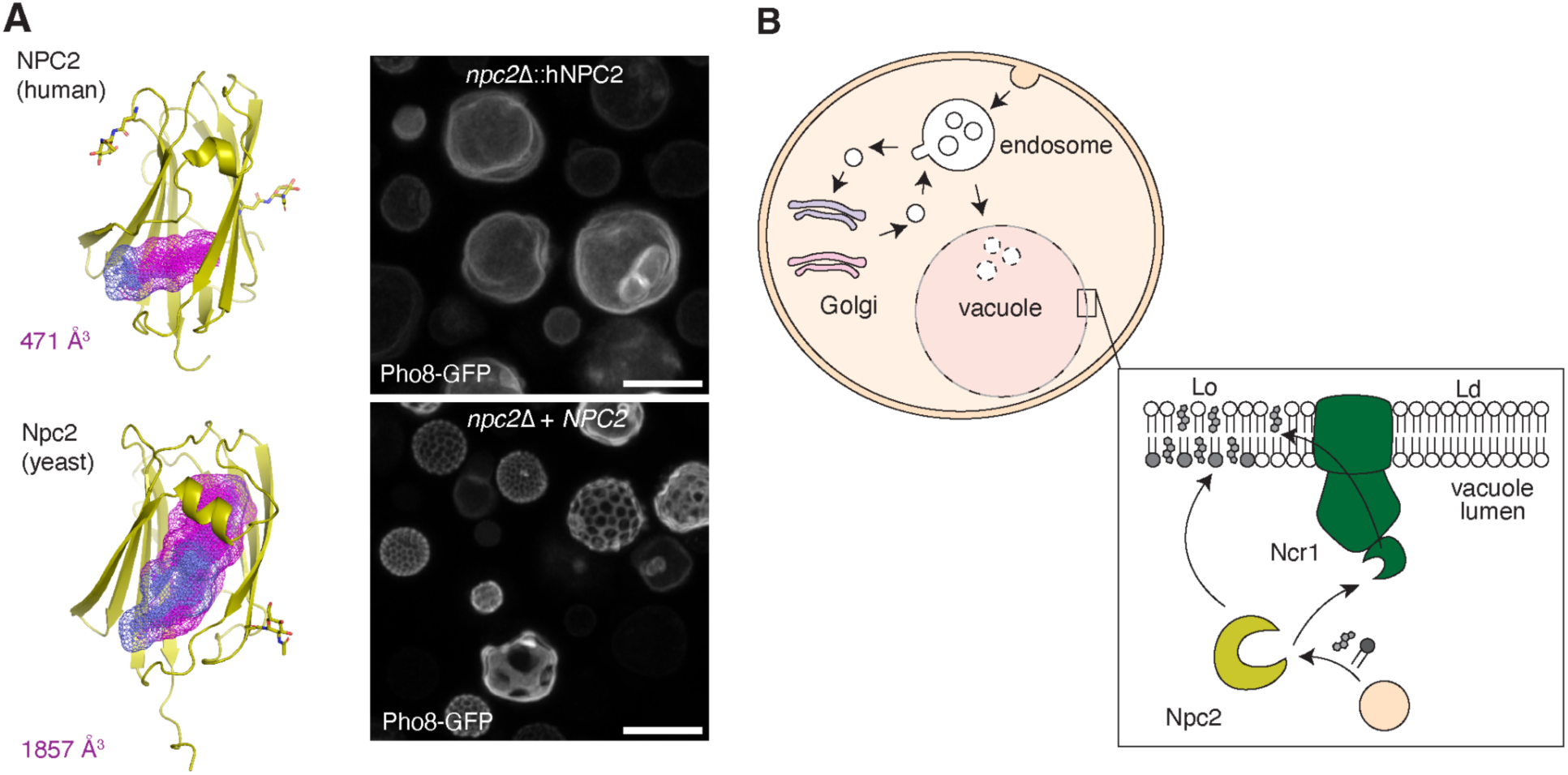
Proposed role for Npc2 in sphingolipid trafficking during vacuole membrane phase separation. **A.** Left: The human NPC2 (top, PDB: 5KWY, chain C) features a smaller hydrophobic binding cavity than the yeast Npc2 (bottom, PDB: 6R4N, chain C). Cavities point clouds were calculated using the Pymol CavitOmiX plugin and cavity sizes were calculated using a modified LIGSITE algorithm (49) as previously described (50). Right: hNPC2 expression fails to rescue the loss of vacuole domains in *npc2*Δ (top), while expression of *NPC2* does. **B.** Model for sphingolipid trafficking to the vacuole membrane. Sphingolipids are endocytosed from the PM, then either recycled to the Golgi apparatus or enter vacuole through the endosomal pathway into the vacuole. Upon late endosome fusion with vacuole, sphingolipids and other lipid cargoes are internalized into the vacuole lumen, where their transport by the NPC system allows incorporation into the vacuole membrane.

## Discussion

Compared to other major lipid classes, transport of sphingolipids between organelles is still not well understood. Previously, we found that sphingolipid trafficking in yeast is required for the phase separation of the vacuole membrane as cells enter stationary stage, as phase-separated vacuoles become enriched in all three yeast complex sphingolipids, as well as ceramides. Here, we leveraged this observation to investigate sphingolipid trafficking to the vacuole under domain forming conditions. Our hypothesis was that trafficking mutants with altered vacuole domains would also show defects in sphingolipid trafficking. We tested this with a BODIPY-tagged ceramide analogue, as well as a previously developed method for isolation of stationary stage vacuoles. We could observe a dependence on both sphingolipid distributions and vacuole domains upon loss of components putatively involved in both intracellular sorting as well as intravacuolar transport. For the latter, we found that incorporation of sphingolipids into the vacuole membrane was dependent on Npc2, but not its canonical receptor Ncr1. This surprising finding led us to test whether yeast Npc2 can serve as an independent transporter of sphingolipids like ceramide and IPC. The identified capacity of Npc2 to transport complex sphingolipids complements recent measurements showing that it can bind a range of lipids, including those derived from native membranes (43).

In mammalian cells, endosomal trafficking is considered a primary route for sphingolipid trafficking into lysosomes (51). In yeast, however, it has not been clear if sphingolipids are similarly trafficked through the endosomal pathway to the vacuole. Upon fusion of the late endosomes with vacuole, intraluminal vesicles are released into the vacuole lumen (52). Similarly, autophagic pathways, which also contribute to domain formation, lead to the release of membrane-bound cargoes into the vacuole lumen. Our results suggest that these diverse cargoes supply sphingolipids for the vacuole membranes. Notably, we still observed robust vacuole staining by BODIPY-Cer after extended incubations in both *vps29*Δ and *vps30*Δ cells, indicating that endosomal trafficking is not the sole source of vacuole sphingolipids. Regardless of their source, lipid transport mechanisms would then be needed to extract complex sphingolipids from intraluminal cargoes, and our results implicate the lipid transporter Npc2 in this process. Such a process would explain the dependence of vacuole domain formation – which corresponds to higher levels of sphingolipids and ergosterol – and subsequent microlipophagy (25) on Npc2 activity.

In the canonical model for NPC system function, Npc2/NPC2 binds its lipid ligand (e.g. cholesterol) and transfers it to the membrane receptor Ncr1/NPC1 via its N-terminus domain (NTD). The ligand is then transported within Ncr1/NPC1 and incorporated into the vacuole/lysosome membrane, where it can be egressed. While NPC2 does transfer sterols to NPC1’s NTD *in vitro* (42), it also can transport them to liposome donors (23, 53). Structures of Ncr1 have provided a potential function for the receptor in allowing transport across the vacuole glycocalyx through an extended sterol transfer tunnel formed by the C-terminal domain (CTD) and middle luminal domain (MLD). This tunnel is of sufficient length to pass the glycocalyx, which has been proposed to pose a physical impediment for Npc2 to directly deposit its ligand into the vacuole membrane (22). Support for a thick vacuole glycocalyx is derived from its similarities to mammalian lysosomes, where NPC1 phenotypes are dependent on glycosylation (54), and the capacity for glycan-binding lectins, like concanavalin A (ConA) to bind to yeast vacuoles *in vitro* (24). However, the exact nature of the vacuole glycocalyx is not resolved, as ConA must be added externally and thus binds to glycans that become cytoplasmically exposed during sample preparation. The inability of Npc2 to interact with glycan modified membranes has also not been established.

Multiple lines of data presented here suggests that Npc2 is capable of direct transport of some lipids to the vacuole membrane, at least under specific conditions in which this process is required for vacuole phase separation. Both vacuole domain formation (Fig. 3A) and the sorting of raft-forming lipids (sphingolipids and sterols) into the vacuole membrane (Fig. 4B) are reduced in *npc2*Δ, but not in *ncr1*Δ, cells. A loss of Npc2, but not Ncr1, also causes intraluminal accumulation of BODIPY-Cer (Fig. 3D). While the Ncr1 NTD can bind to larger lipids compared to its mammalian homologues (42), it is not clear if sphingolipids with large polar head groups, like IPC, could be accommodated by the narrow and hydrophobic binding tunnel of Ncr1. Our results indicate that Npc2 can transport sphingolipid analogues (e.g. NBD-IPC) between membranes more robustly than it does sterols.

There are several potential models to explain an Ncr1-independent role for Npc2 function in vacuole domain formation. The glycocalyx could not pose a sufficient barrier to prevent direct vacuole membrane-Npc2 contacts or Npc2 could interact with other, unidentified receptors in *ncr1*Δ cells. An alternative possibility is that the glycocalyx is no longer uniformly distributed in phase separated stationary stage vacuoles, and this can promote additional incorporation of Lo-domain promoting lipids. We observed that some isolated vacuoles show congruence between ConA and the Ld marker, Vph1-mCherry (Figure S8), suggesting that luminal glycans might themselves laterally organize separately alongside lipids and proteins. A major limitation of these experiments remains inconsistent ConA staining of vacuoles, presumably due to the need for glycans to become cytoplasmically exposed. New methods to measure luminal glycans and their interactions within lipid transporters are needed to fully evaluate these hypotheses.

## Acknowledgements

Alessandro Fracassi, Caroline Knittel, Zulfiqar Mohamedshahand, and Neal Devaraj provided instrumentation and technical assistance for compound synthesis and characterization. Yongxuan Su and the UCSD Molecular Mass Spectrometry Facility aided in protein analysis. Xuemei Huang and UCSD Biomolecular NMR facility aided in NMR analysis. Arnold Seo provided reagents and technical support. Research was supported by grants from the National Institutes of Health (NIH) (R35-GM142960) and the National Science Foundation (NSF) (MCB-2046303) to I.B. I.J-C. was supported by the NIH Molecular Biophysics Training Grant (T32-GM008326C). A.M. W. was supported by the NSF Graduate Research Fellowship Program (DGE-2038238) and the NIH Chemistry and Biology Interface Training Grant (T32-GM146648).

## Supporting Information

**Table S1.**
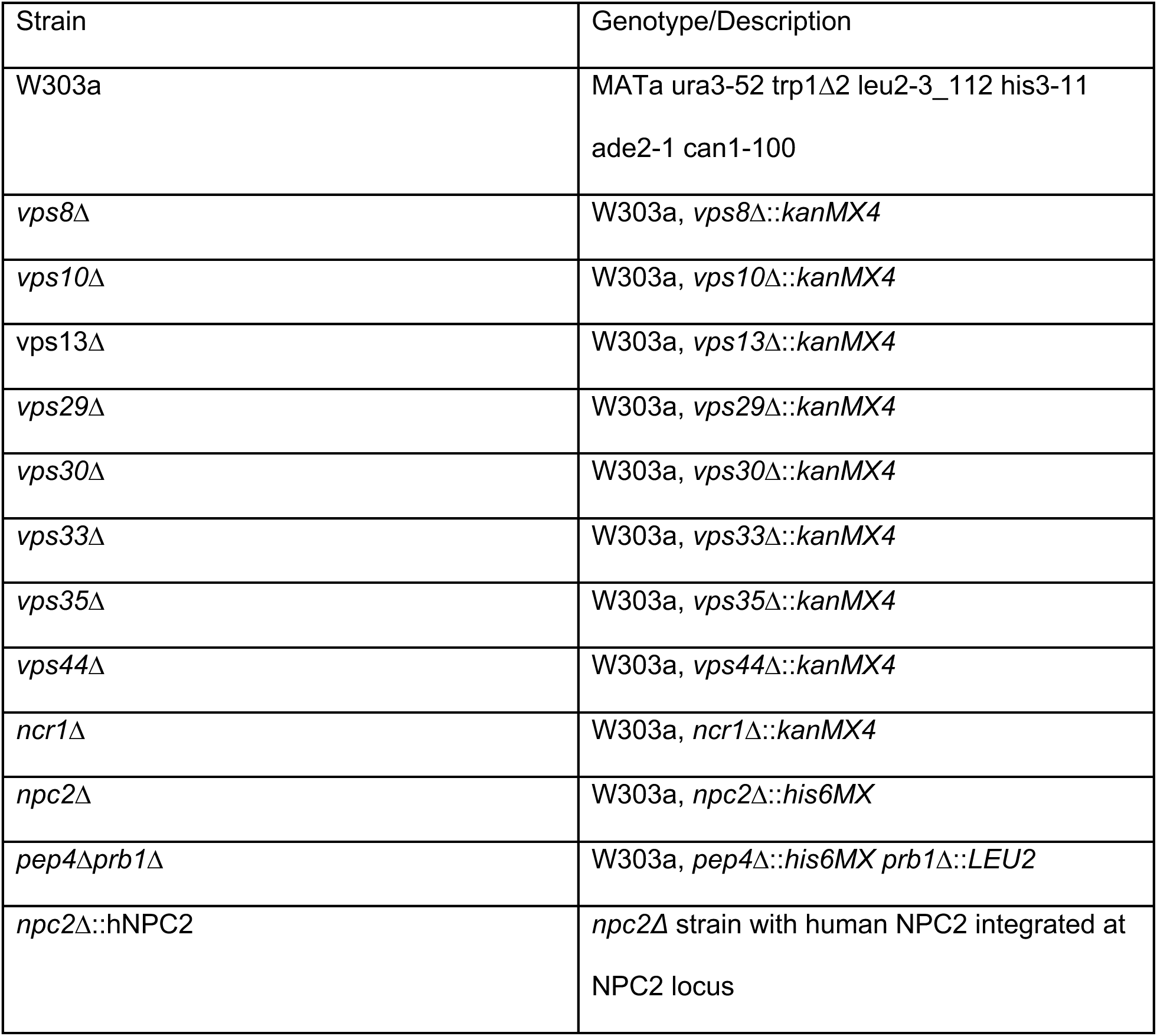
List of yeast strains used in this study.

| Strain | Genotype/Description |
| --- | --- |
| W303a | MATa ura3-52 trp1Δ2 leu2-3_112 his3-11 ade2-1 can1-100 |
| *vps8*Δ | W303a, *vps8*Δ::*kanMX4* |
| *vps10*Δ | W303a, *vps10*Δ::*kanMX4* |
| *vps13*Δ | W303a, *vps13*Δ::*kanMX4* |
| *vps29*Δ | W303a, *vps29*Δ::*kanMX4* |
| *vps30*Δ | W303a, *vps30*Δ::*kanMX4* |
| *vps33*Δ | W303a, *vps33*Δ::*kanMX4* |
| *vps35*Δ | W303a, *vps35*Δ::*kanMX4* |
| *vps44*Δ | W303a, *vps44*Δ::*kanMX4* |
| *ncr1*Δ | W303a, *ncr1*Δ::*kanMX4* |
| *npc2*Δ | W303a, *npc2*Δ::*his6MX* |
| *pep4*Δ*prb1*Δ | W303a, *pep4*Δ::*his6MX* *prb1*Δ::*LEU2* |
| *npc2*Δ::hNPC2 | *npc2*Δ strain with human NPC2 integrated at NPC2 locus |

**Figure S1.**
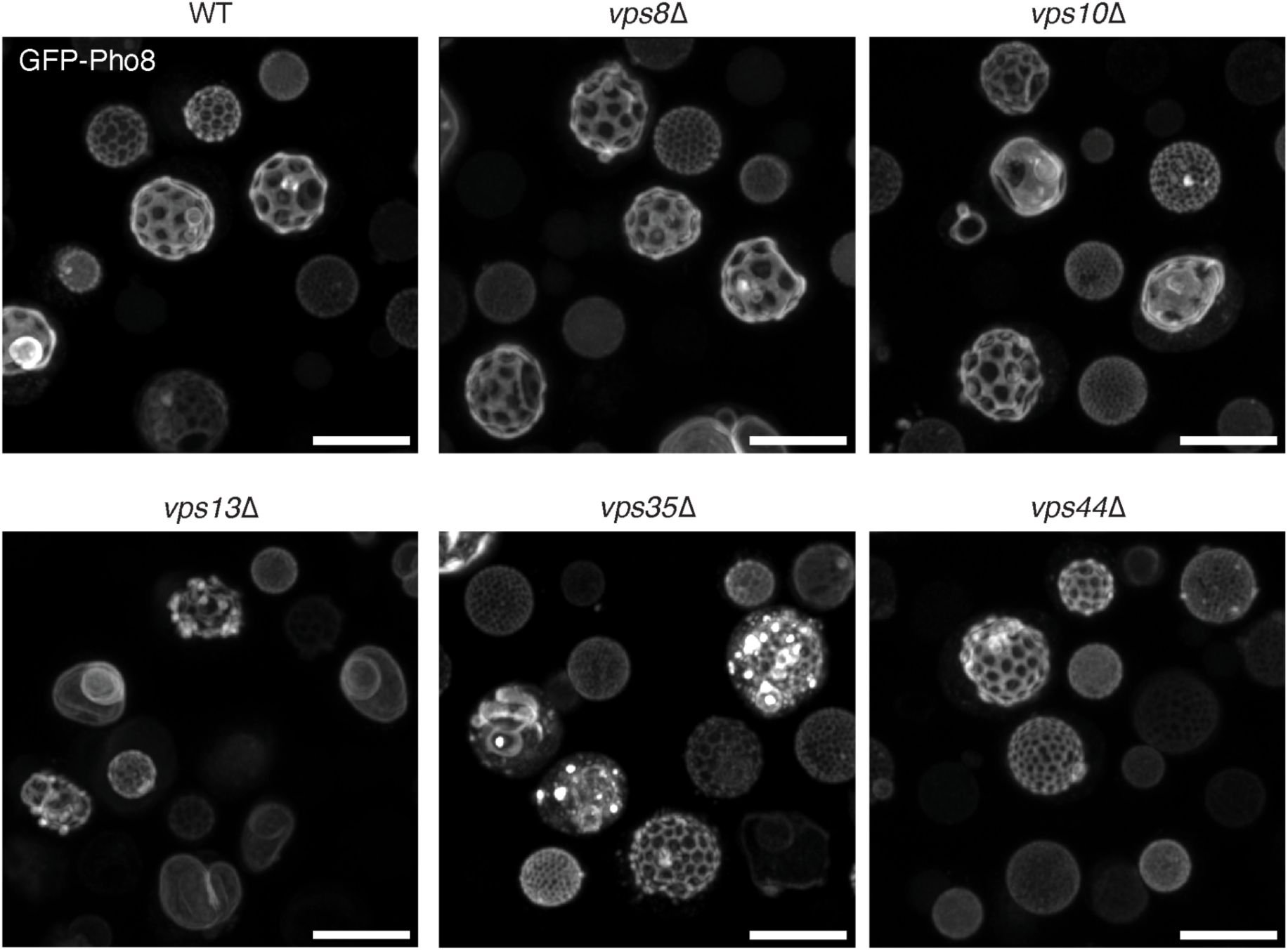
Images of yeast vacuole domains of other Type A mutants. Scale bar, 5µm.

**Figure S2.**
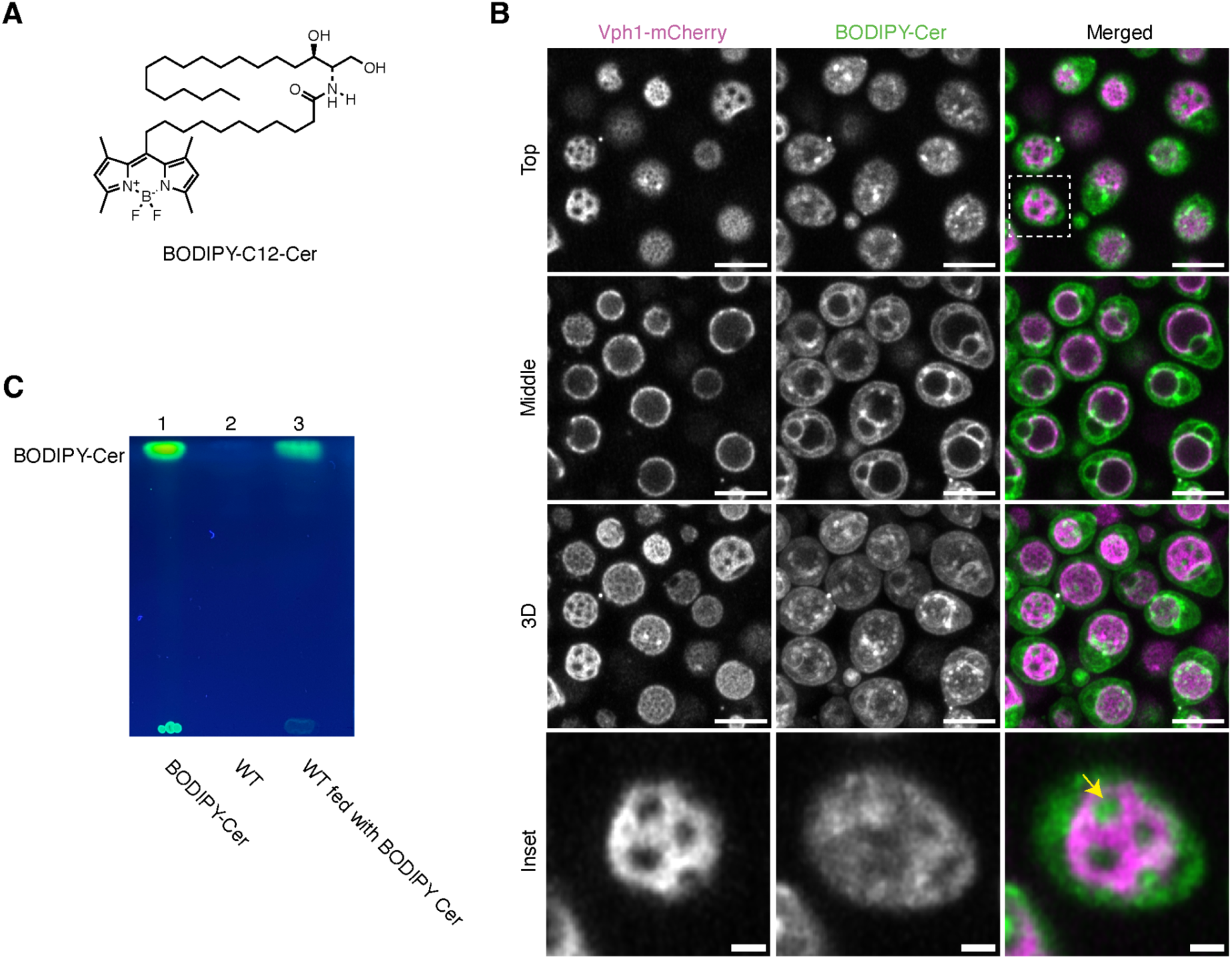
BODIPY-Cer is not converted to complex sphingolipids in WT cells. **A.** chemical structure of BODIPY-C12-Cer used in this study. **B.** Micrographs of WT cells expressing Vph1-mCherry and fed with BODIPY-Cer. The Inset from the top view shows partial localization of BODIPY-Cer in Lo domain, indicated by the arrow. Scale bar, 5 µm for Top, Mid and 3D view. Scale bar, 1 µm for the inset. **C.** TLC of sphingolipid extracted from WT with or without BODIPY-Cer, which were run alongside a BODIPY-Cer stock solution. The TLC plate was visualized under UV. No additional fluorescent bands were detected in cell extracts, indicating no further conversion of BODIPY-Cer to complex sphingolipids.

**Figure S3.**
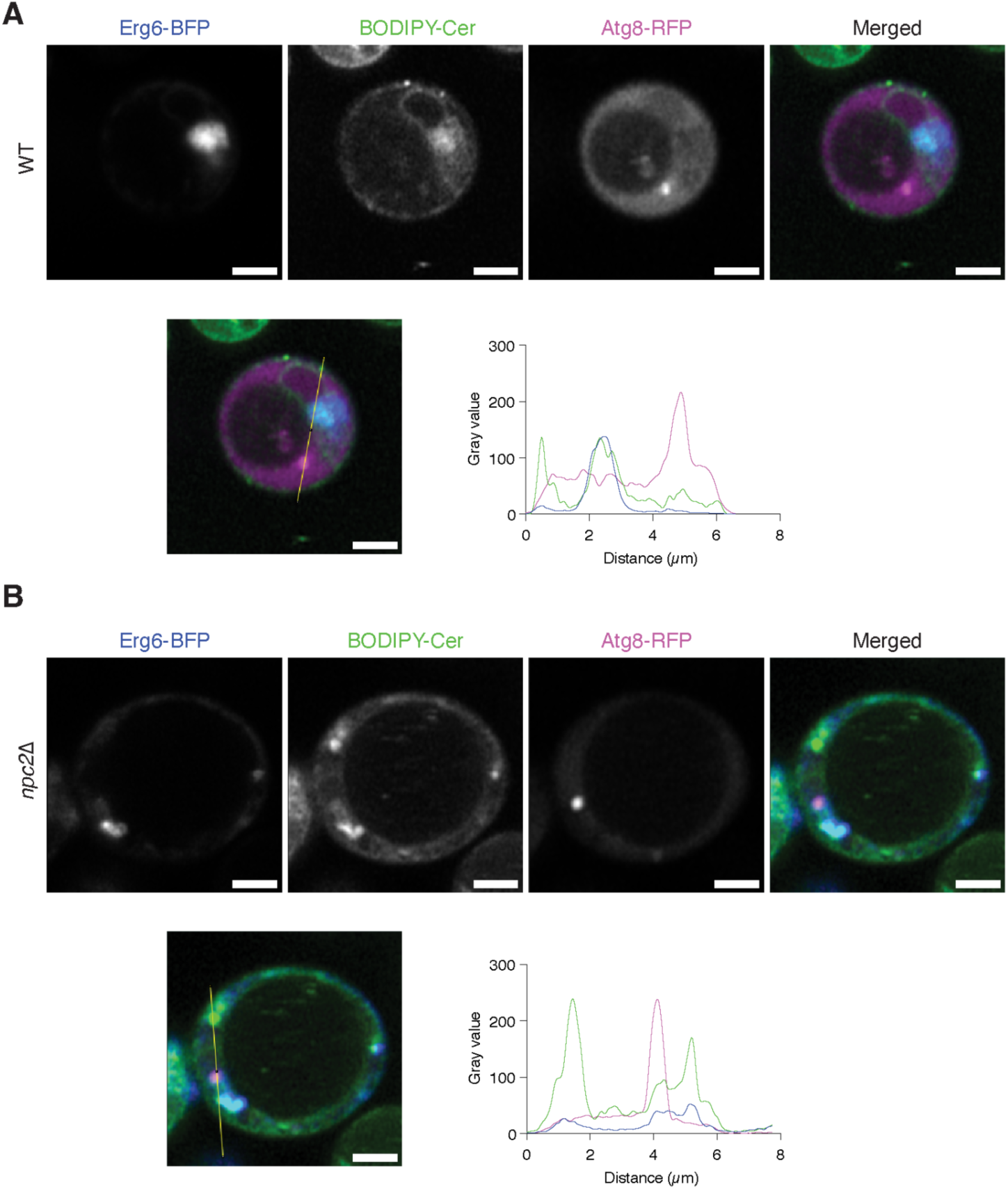
BODIPY-Cer signal partially colocalizes with the lipid droplet marker, Erg6-BFP, and does not with autophagosome marker Atg8-RFP in (**A**) WT and (**B**) *npc2*Δ cells. Erg6-BFP and Atg8-RFP were expressed by plasmid transformation and the cells were stained with BODIPY-Cer for 1 hour. The line profile was taken across different colored puncta to determine whether the BODIPY-Cer puncta colocalizes with lipid droplet or autophagosome. Scale bar, 1 µm.

**Figure S4.**
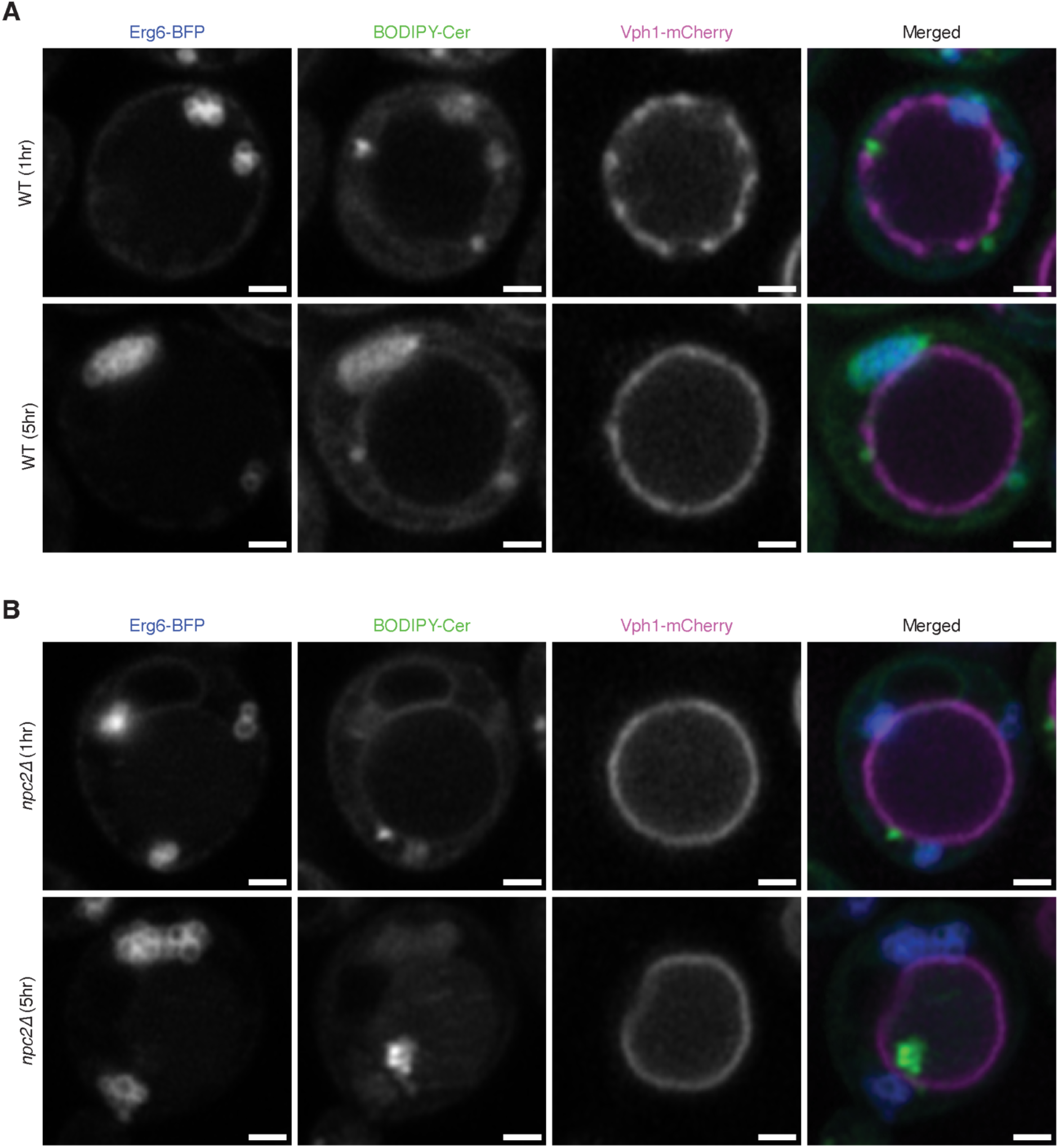
BODIPY-Cer punctae at or inside the vacuole are not lipid droplets. **A.** The small punctae of BODIPY-Cer found peripheral to the vacuole in WT cells do not colocalize with Erg6-BFP. Scale bar, 1µm. **B.** In cells lacking Npc2, larger aggregates of BODIPY-Cer accumulate within the vacuole over extended incubations. These do not show co-localization with Erg6-BFP, indicating that they are endosomal in origin. Scale bar, 1µm.

**Figure S5.**
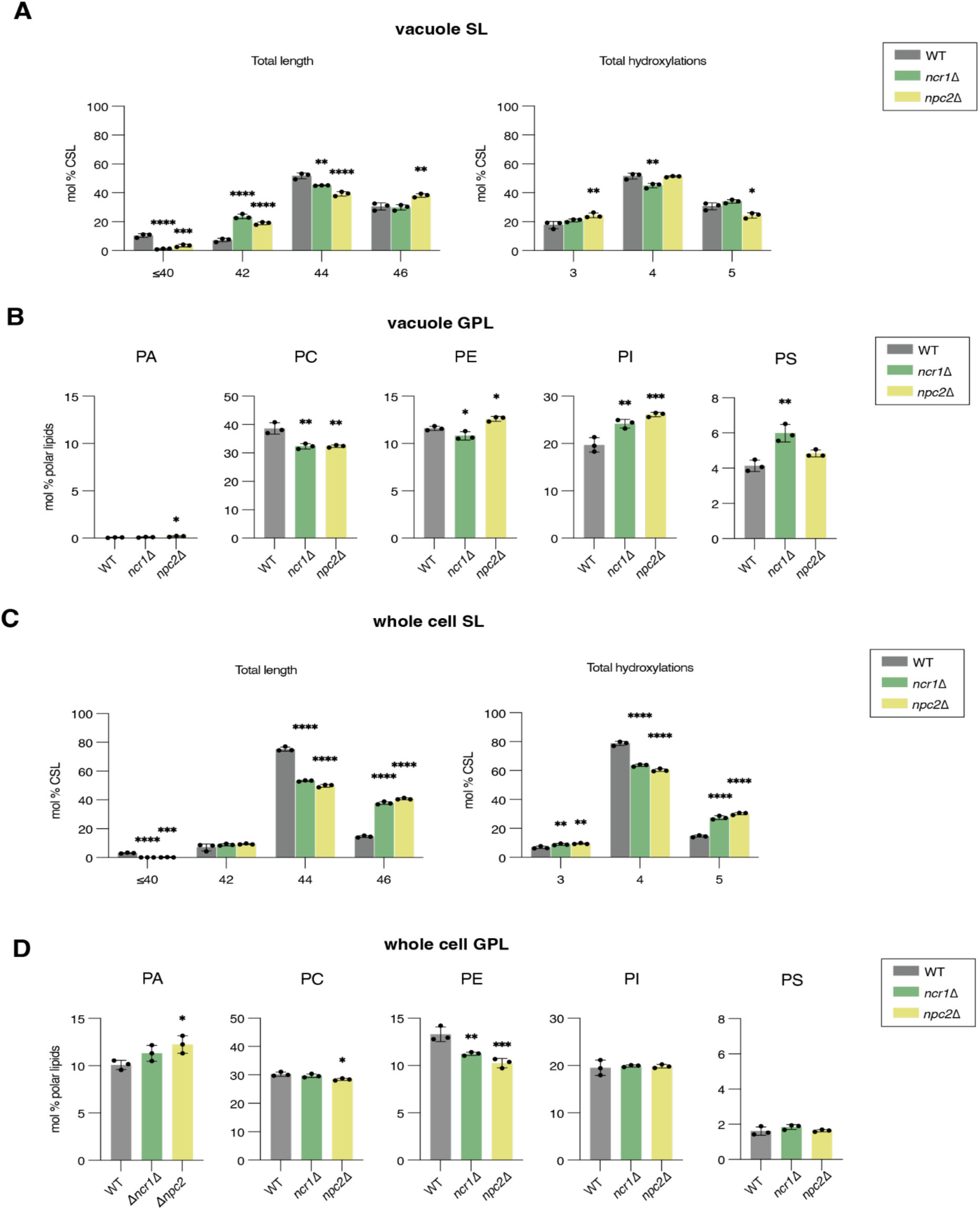
Changes in glycerophospholipid (GPL) and sphingolipid (SL) lipidome of *ncr1*Δ and *npc2*Δ early stationary cells and vacuoles. **A.** Changes in SL composition by length of acyl chain and hydroxylation in the vacuole. **B.** Vacuolar GPL composition by phospholipid class. The low PA levels are characteristic of purified vacuoles. **C.** Whole cell SL composition by length of acyl chain and hydroxylation state. **D.** Whole cell GPL composition grouped by phospholipid class. Phosphatidylglycerol (PG) is not shown due to its low abundance in the cell. For all panels, significance against WT was assessed one-way ANOVA with Dunnett’s post-hoc test; *p < 0.05; **p < 0.01; ***p < 0.001, ****p < 0.0001.

**Figure S6.**
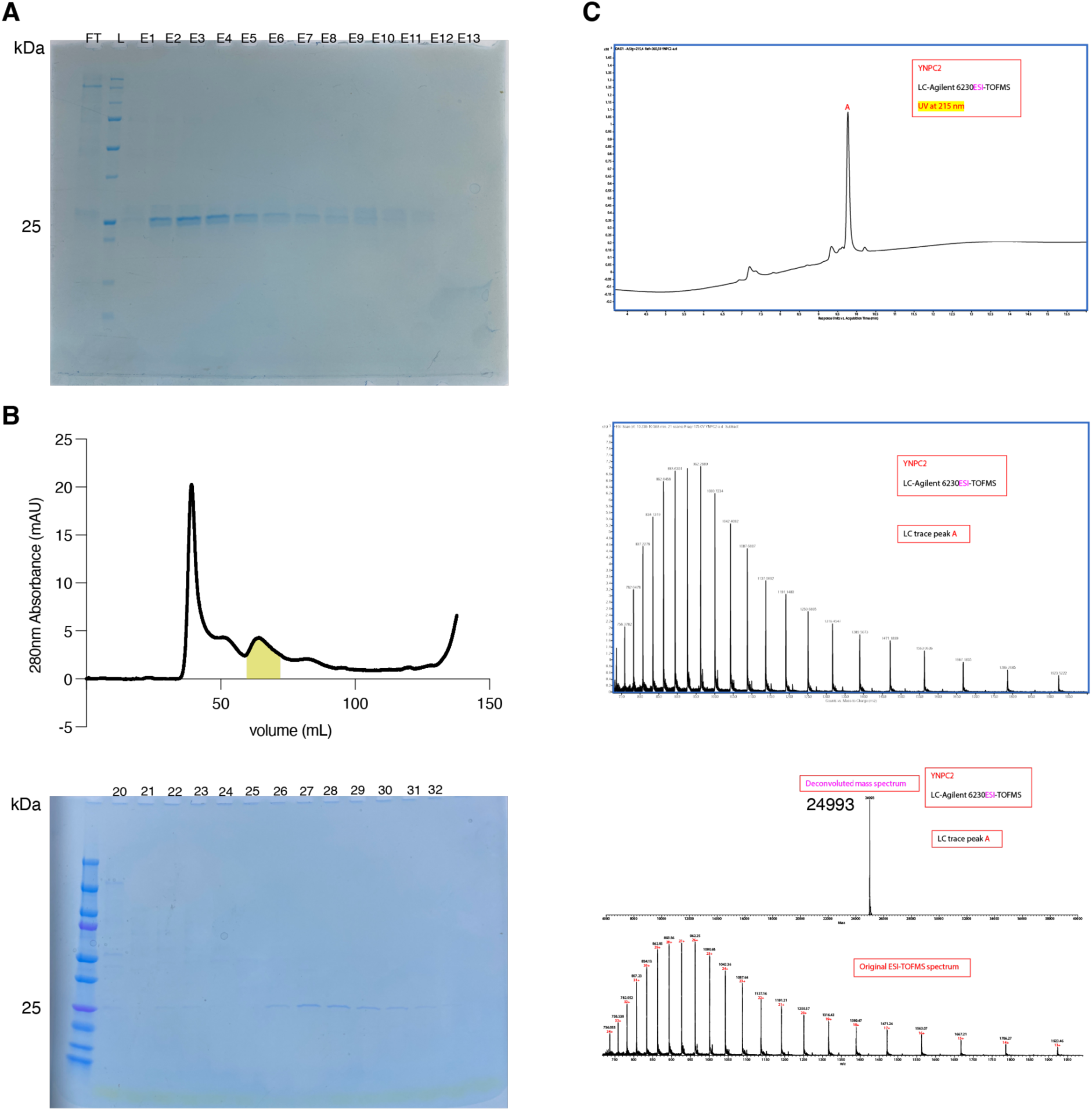
Purification and characterization of Npc2 from yeast cells. **A.** SDS-PAGE gel following His-Tag affinity chromatography. FT, flow-through; L, protein ladder; E, elution. **B.** Size-exclusion chromatogram (top) with corresponding SDS-PAGE of the highlighted 2.0 mL fractions (bottom). **C.** Mass spectrometry of purified Npc2. The theoretical molecular weight of the protein is 21.67 kDa, including the 10x His and thrombin tag, but the observed molecular weight of purified protein is 24.99 kDa due to its glycosylation.

**Figure S7.**
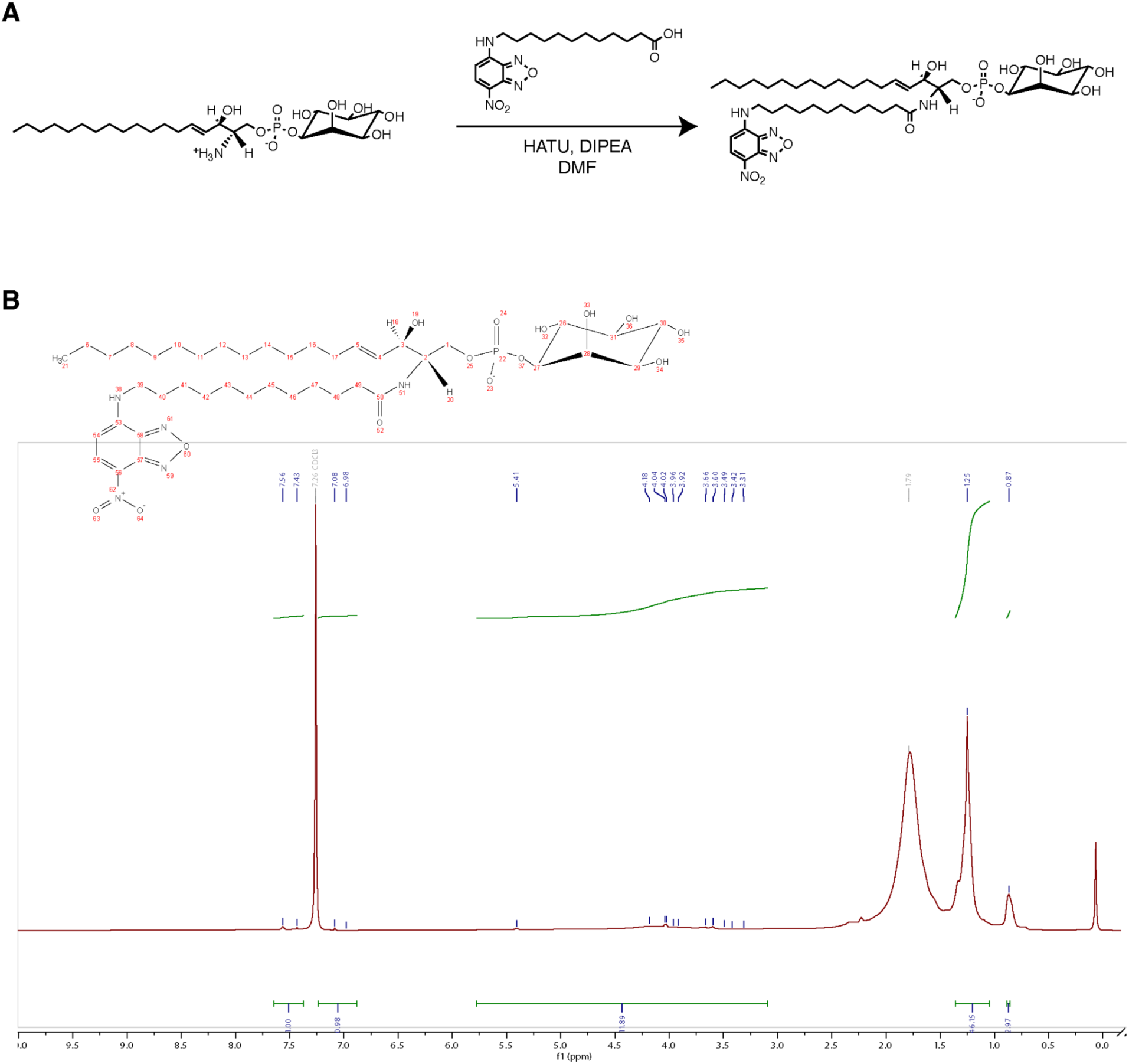
Synthesis and characterization of NBD-IPC. **A.** Reaction scheme showing the HATU coupling of sphingosyl-PI and C-12 NBD. **B.** ^1^H-NMR spectrum of NBD-IPC and selected peaks integrated for analysis.

**Figure S8.**
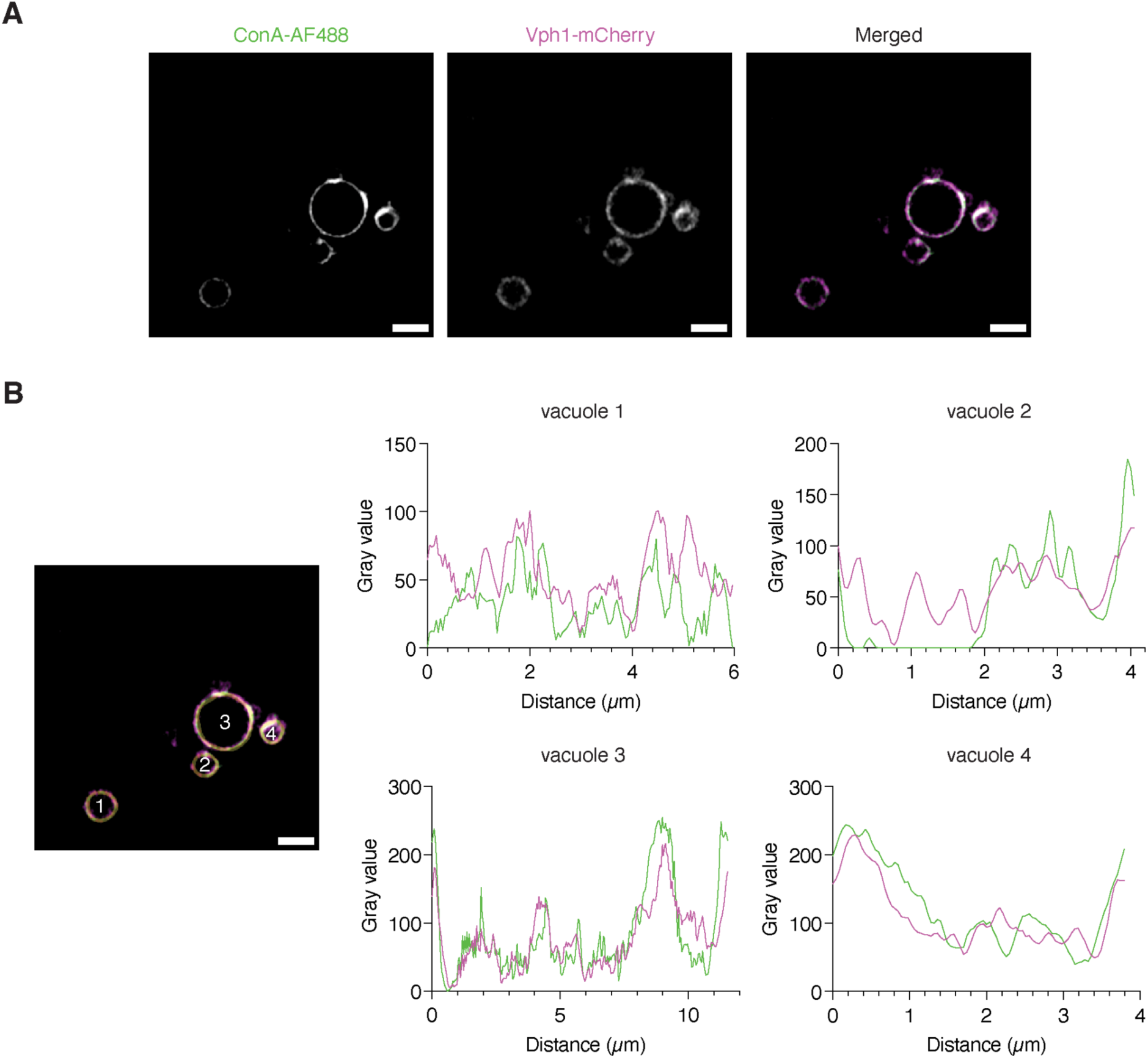
Co-localization of vacuole glycan patches with the Ld marker Vph1-mCherry. **A.** Purified vacuoles from cells expressing Vph1-mCherry and stained with ConA conjugated AlexaFluor 488 (ConA-AF488). Scale bar, 2µm. **B.** Line profiles showing variations in both Ld and glycan markers along vacuole perimeters. Some (vacuole 3) show complete co-localization, while others (1, 2, 4) show incomplete localization or co-localization only in patches.

**Data File S1.** Spreadsheet reporting the abundances of each lipid species, expressed in pmol per sample, in the whole cell and vacuole samples used in this study. For each sample, three biological replicates are provided. Mol % values were calculated by dividing the abundance of each individual lipid species by the abundances summed for all species in the same sample. For calculation of mol % of polar lipids, TAGs, DAGs, and EEs were excluded.

